# Reconstruction of motor control circuits in adult *Drosophila* using automated transmission electron microscopy

**DOI:** 10.1101/2020.01.10.902478

**Authors:** Jasper T. Maniates-Selvin, David Grant Colburn Hildebrand, Brett J. Graham, Aaron T. Kuan, Logan A. Thomas, Tri Nguyen, Julia Buhmann, Anthony W. Azevedo, Brendan L. Shanny, Jan Funke, John C. Tuthill, Wei-Chung Allen Lee

## Abstract

Many animals use coordinated limb movements to interact with and navigate through the environment. To investigate circuit mechanisms underlying locomotor behavior, we used serial-section electron microscopy (EM) to map synaptic connectivity within a neuronal network that controls limb movements. We present a synapse-resolution EM dataset containing the ventral nerve cord (VNC) of an adult female *Drosophila melanogaster*. To generate this dataset, we developed GridTape, a technology that combines automated serial-section collection with automated high-throughput transmission EM. Using this dataset, we reconstructed 507 motor neurons, including all those that control the legs and wings. We show that a specific class of leg sensory neurons directly synapse onto the largest-caliber motor neuron axons on both sides of the body, representing a unique feedback pathway for fast limb control. We provide open access to the dataset and reconstructions registered to a standard atlas to permit matching of cells between EM and light microscopy data. We also provide GridTape instrumentation designs and software to make large-scale EM data acquisition more accessible and affordable to the scientific community.

## INTRODUCTION

To navigate a complex world, an animal’s nervous system must stimulate precise patterns of muscle contractions to give rise to coordinated body movements. Humans have more than 600 skeletal muscles containing 100 million muscle fibers, which are innervated by more than 100,000 motor neurons (Kanning et al., 2010). Limb motor neurons reside in the spinal cord, where neuronal networks integrate signals from the brain with sensory feedback from the body to coordinate limb movements. A century of studies in mammals has revealed many principles of spinal cord organization and development (Kiehn, 2011). However, we still lack detailed knowledge of the wiring and connectivity patterns of neuronal circuits that control motor output.

Insects have a ventral nerve cord (VNC) that is homologous to the vertebrate spinal cord (Niven et al., 2008), but insects lack vertebrae, making the VNC more experimentally accessible. Moreover, insects also have complex locomotor behaviors, and many of their neurons are uniquely identifiable across different individuals. As a result, insects are well-established models for understanding the physiology of motor neurons and premotor circuits, which reside in the VNC (Burrows, 1996; Buschges et al., 2008).

*Drosophila melanogaster* is a particularly appealing system for dissecting mechanisms of motor control because it is a genetically accessible model system and has many complex and well-characterized behaviors including walking, flight, escape responses, grooming, and courtship. Recent advances allow *in vivo* electrophysiological recordings and calcium imaging of genetically identified VNC neurons in behaving adult *Drosophila* (Chen et al., 2018; Mamiya et al., 2018; Tuthill and Wilson, 2016b), providing an increasingly better grasp of motor and premotor neuron activity during locomotor behavior. Furthermore, the small size of the *Drosophila* nervous system makes it suitable for comprehensive connectome mapping using electron microscopy (EM). Connectomic reconstruction was previously undertaken for a larval *Drosophila* nervous system (Ohyama et al., 2015; Schneider-Mizell et al., 2016) and an adult brain (Takemura et al., 2013; Tobin et al., 2017; Zheng et al., 2018), but not yet for an adult VNC, which accounts for a third of the adult central nervous system and contains all limb motor neurons. A VNC connectome would enhance our understanding of how VNC circuits control muscles of the legs (Soler et al., 2004), neck (Strausfeld et al., 1987), wings (O’Sullivan et al., 2018), and halteres (Dickerson et al., 2019) to give rise to complex behaviors.

Each of *Drosophila’s* six legs is controlled by 14 intrinsic muscles and ∼53 motor neurons (Baek and Mann, 2009; Miller, 1950; Soler et al., 2004). From each motor neuron’s cell body arises a single process called a primary neurite which travels through the VNC and then exits through a peripheral nerve, at which point the primary neurite becomes an axon that carries action potentials to muscles (Burrows, 1996). When traveling through the VNC, the primary neurite gives rise to numerous thinner branches onto which the motor neuron receives synaptic input. Most motor neurons have branching patterns that are morphologically distinct (Baek and Mann, 2009; Brierley et al., 2012) and many are genetically identifiable (Enriquez et al., 2015; Venkatasubramanian et al., 2019). Motor neurons controlling the legs receive synaptic input within one of the VNC’s six thoracic neuromeres, roughly spherical compartments that together comprise most of the VNC. Each neuromere corresponds primarily to a single leg. The front, middle, and hind legs and their corresponding neuromeres are termed T1, T2, and T3, respectively. Motor neuron dendrites occupy specific regions of the neuromere, with motor neurons that control muscles in the same leg segment typically occupying the same regions of the neuromere (Baek and Mann, 2009; Brierley et al., 2012). In addition to their morphological diversity, motor neurons have a gradient of anatomical, physiological, and functional properties. “Fast” motor neurons control ballistic, large-amplitude movements, “slow” neurons control postural, small-amplitude movements, and other motor neurons fall between these two extremes (Azevedo et al., 2019).

Flexible motor control relies heavily on sensory feedback from proprioceptors, a class of sensory neurons that measure body position, velocity, and load. In both vertebrates and invertebrates, proprioceptive feedback is processed by the central nervous system to fine-tune motor output (Tuthill and Azim, 2018). The major proprioceptor types in insects are chordotonal organs, hair plates, and campaniform sensilla, each innervated by sensory neurons with distinct structural and functional properties (Tuthill and Wilson, 2016a). Morphologically distinct subclasses of chordotonal neurons encode different features of leg movement such as position, velocity, and vibration (Mamiya et al., 2018). Campaniform sensilla encode load signals similar to mammalian Golgi tendon organs (Pringle, 1938; Tuthill and Azim, 2018; Zill and Moran, 1981). Although we know about basic proprioceptor types and the signals they encode, we lack an understanding of how motor circuits integrate proprioceptive inputs to control the body.

Electron microscopy (EM) is the gold standard for mapping structural connectivity within neuronal circuits (Sjostrand, 1958; White et al., 1986). However, even seemingly small tissue volumes (1 mm^3^) acquired at synaptic resolution (e.g. 4 × 4 × 40 nm^3^ per voxel) produce massive datasets (>1500 teravoxels) that require automated methods for reliable acquisition in a reasonable amount of time. Recent developments in scanning electron microscopy (SEM) methods enabled connectomic analyses of multiple neuronal circuits (Briggman et al., 2011; Hildebrand et al., 2017; Kasthuri et al., 2015; Kornfeld et al., 2017; Morgan et al., 2016; Schmidt et al., 2017; Tapia et al., 2012; Wanner et al., 2016). Compared to SEM, transmission EM (TEM) allows for higher spatial resolution (Merk et al., 2016), an order of magnitude greater signal-to-noise at the same electron dose (Xu et al., 2017; Zheng et al., 2018), and straightforward parallelization (Bock et al., 2011; Lee et al., 2016; Tobin et al., 2017; Zheng et al., 2018). Although there have been recent developments in motorized TEM section collection (Lee et al., 2018) and automated high-throughput TEM imaging (Zheng et al., 2018), we lack an end-to-end platform for automated high-throughput TEM section collection and imaging. To address this, we designed a tape-based data acquisition pipeline that combines automated sectioning with a novel TEM-compatible collection substrate and an automated, reel-to-reel imaging stage. This technology, called GridTape, accelerates section collection and enables sustained TEM imaging rates of >40 Mpixels per second for a fraction of the cost of alternative systems.

Here, we used GridTape to produce a synapse-resolution EM dataset of the VNC of an adult female *Drosophila melanogaster*. We reconstructed sensory and motor neurons to reveal a neuronal network that controls limb movements. We found that sensory and motor axons occupy specific spatial domains within leg nerves. We identified a new class of leg proprioceptive neuron, the bilaterally projecting campaniform sensillum (bCS), that projects to multiple neuromeres to provide direct synaptic input onto the motor neurons with the largest-diameter axons associated with each leg. We registered the EM dataset to a light microscopy–based VNC atlas, allowing us to identify a functionally characterized “fast” flexor motor neuron as strong synaptic target of bCS neurons. Our reconstructions also demonstrated that many motor neurons have branching patterns that are morphologically unique and bilaterally symmetric. We provide the EM dataset, neuron reconstructions, and designs for GridTape instrumentation as freely available resources for the scientific community.

## RESULTS

### GridTape: an accessible TEM platform for connectomics

We developed GridTape, a TEM-compatible tape substrate (Fig. 1A) that combines advantages of automated section collection from the automated tape-collecting ultramicrotome SEM (ATUM-SEM) approach (Hayworth et al., 2014) with the advantages of TEM imaging (Figs. 1B-C and S1). To produce GridTape, regularly spaced 2 mm × 1.5 mm holes resembling slots in conventional TEM grids are laser-milled through aluminum-coated polyimide (Kapton®) tape (Fig. 1A). The milled tape is then coated with a 50 nm-thick film (LUXFilm®) that spans the slots to provide support for subsequent section collection and can be safely layered upon itself. We collect sections onto GridTape using an ATUM modified for compatibility with GridTape (Fig. S1A-E). The tape is positioned near the ultramicrotome’s cutting knife so that sections consistently adhere to the moving tape as they are being cut. By monitoring the ultramicrotome cutting speed and adjusting the speed of the tape, the movement of GridTape slots is locked in-phase with cutting. This closed-loop sectioning approach permits automated collection of >4000 sections per day with reliable positioning of sections over film-coated slots.

**Figure 1.**
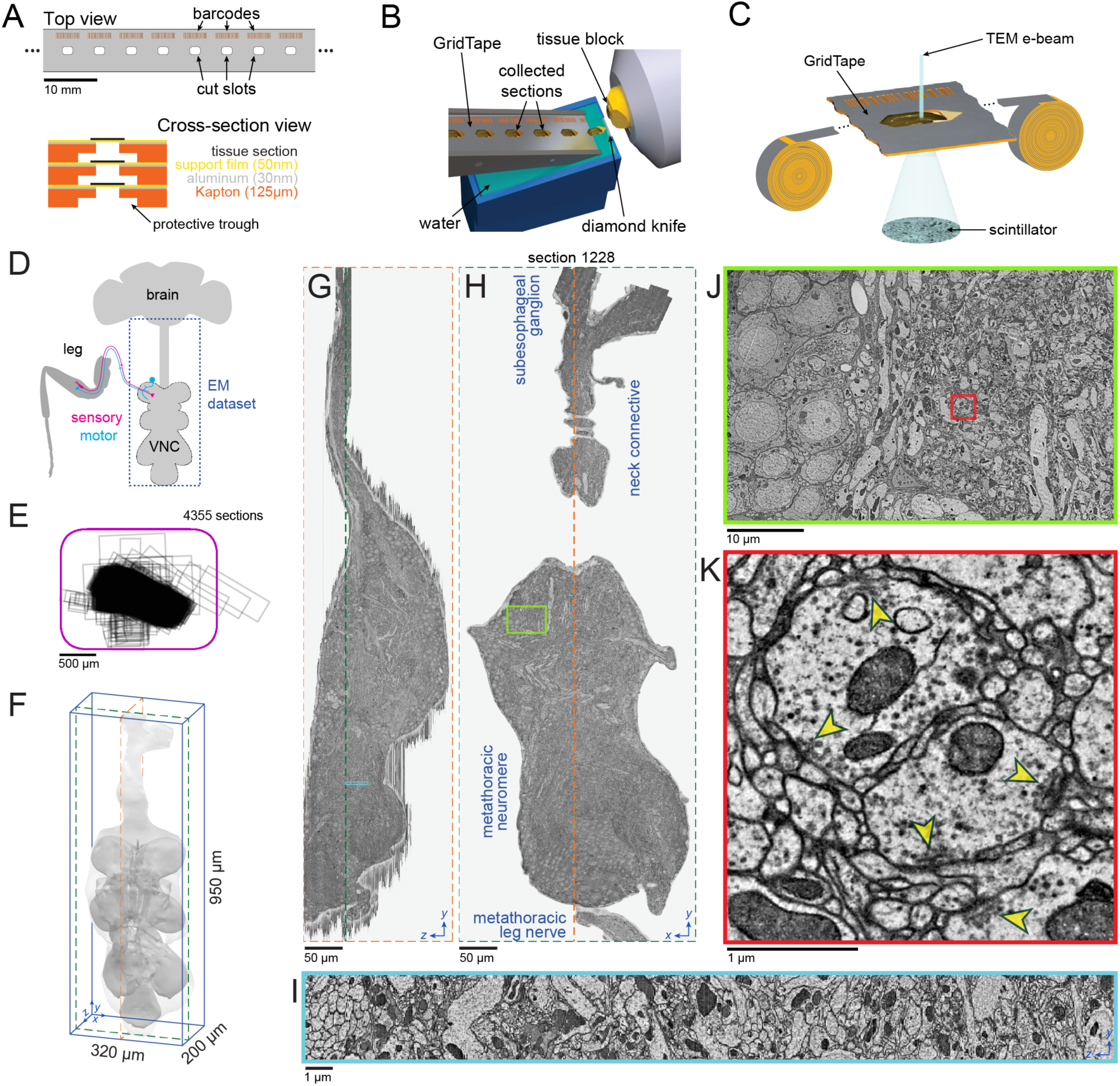
Acquiring a ventral nerve cord (VNC) dataset using a high-throughput serial-section TEM pipeline built around GridTape. (A) (top) Regularly-spaced holes and barcodes are laser-milled into a length of tape to produce GridTape, a substrate for collection of serial sections. (bottom) Schematic of the GridTape layered in cross-section. For clarity, tape thickness is exaggerated. (B) Resin-embedded samples are sectioned using a GridTape-compatible automated tape-collecting ultramicrotome. Sections adhere to GridTape immediately after being cut and are targeted to land over a hole in the tape. See also Fig. S1. (C) Reels of GridTape are inserted into a custom stage attached to a transmission electron microscope, enabling sections to be positioned under the electron beam. See also Fig. S1G. (D) Schematic of the adult Drosophila central nervous system and leg. The synapse-resolution dataset presented here contains the VNC and its connection to the brain (dashed outline). (E) The VNC was cut into 4355 thin sections (∼45 nm) and collected onto GridTape. Each black box indicates the bounding box of the VNC in a single section relative to that section’s slot (purple outline). Two sections were collected off-slot and are not shown. (F) Volumetric rendering of the VNC dataset and measurements of its bounding box. Light grey, the outline of all imaged tissue. Dark grey, the outline of the VNC’s neuropil. (G) A yz-reslice through the aligned dataset at the level of the orange dashed plane in (F) and (H). (H) A single section at the level of the green dashed plane in (F) and (G). The imaged region spans from the suboesophageal ganglion in the ventral brain, across the neck connective to the metathoracic neuromere and the metathoracic leg nerve. (I) Zoomed-in yz-reslice from the region (cyan box) in (G). (Legend continued on next page) (J) Zoom-in from the region (red box) in (H). (K) Zoomed-in view of synaptic contacts from the region (green box) in (J). Yellow arrowheads indicate presynaptic specializations known as T-bars. Scale bars, 10 mm (A, top), 500 µm (E), 50 µm (G-H), 10 µm (J), 1 µm (I, K).

Collecting sections onto thin films enables widefield TEM imaging. To automate the imaging process, we engineered a stage that attaches to standard TEM microscopes and houses reels of GridTape in vacuum (Fig. S1G). Tape housings were added on opposite sides of the microscope column to allow motors to feed the tape between the two sides and position sections under the electron beam for imaging. To image large areas at synaptic resolution, the microscope automatically montages each section using piezoelectric nano-positioners. After each section is imaged, the tape is translated to position the next section for montaging, enabling continuous unattended operation. Using a 2×2 camera array (Bock et al., 2011), we achieve effective imaging rates of >40 Mpixels per second (see Table 1). This microscope, termed TEMCA-GT (Transmission Electron Microscope with a Camera Array and GridTape), provides high-throughput imaging at a relatively low cost of ∼US%300,000 per microscope (Tables 1 and S1).

**Table 1.**
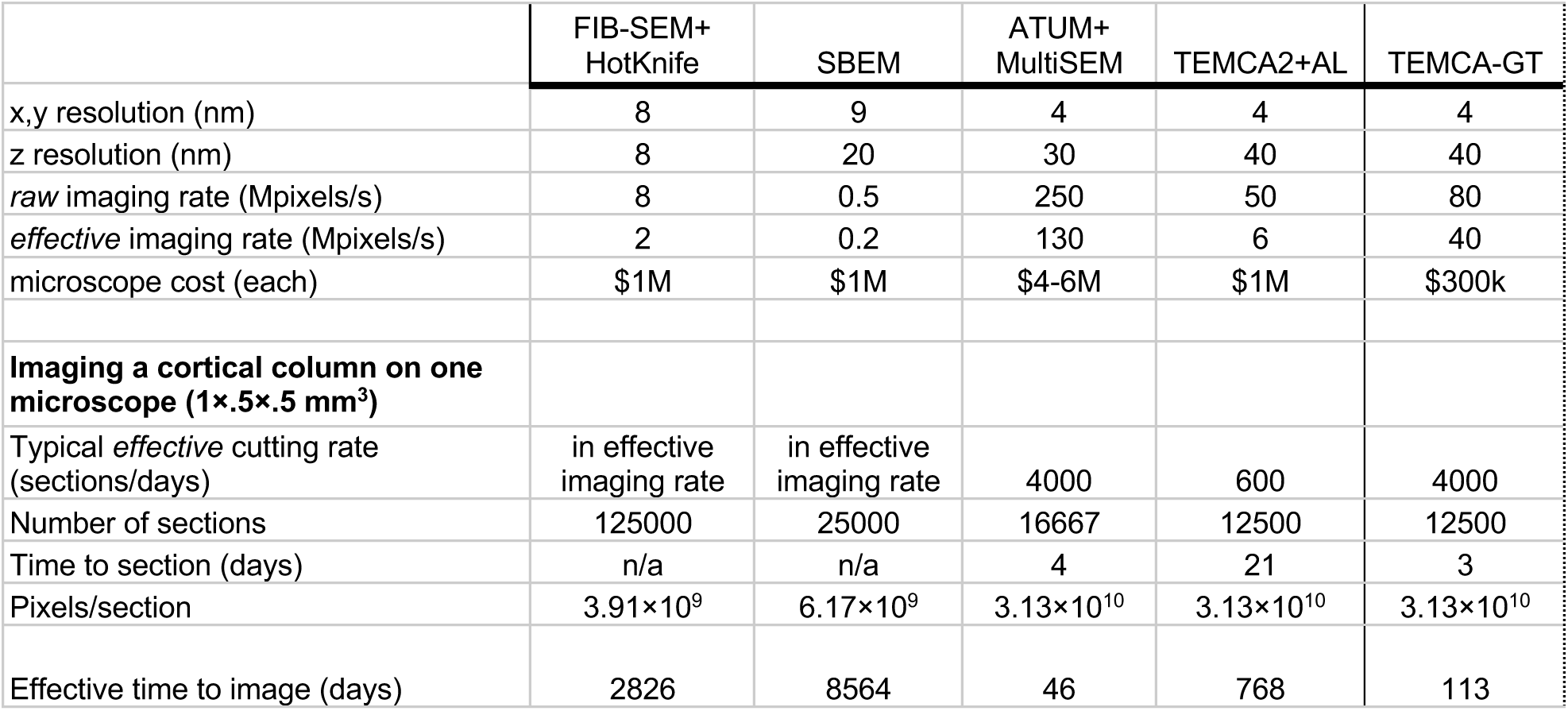
Serial EM microscope throughput and cost comparison. Based on published datasets: focused ion beam milling SEM (FIB-SEM) resolution range: x,y,z: 5–8nm (Knott et al., 2008; Xu et al., 2017), serial block-face SEM (SBEM): resolution ranges x,y: 9–16nm, z: 20–30nm (Briggman et al., 2011; Kornfeld et al., 2017; Schmidt et al., 2017). For FIB-SEM+HotKnife (Hayworth et al., 2015) imaging rates, Ken Hayworth, personal communication; automated tape-collecting ultramicrotome multi-beam SEM (ATUM-MultiSEM) resolution and imaging rates, Richard Schalek, personal communication; and for TEMCA2-AutoLoader (Zheng et al., 2018), Cam Robinson, personal communication. Effective imaging rate is defined as the dataset size divided by calendar days from start to end of imaging, including overhead time such as stage movement, microscope downtime, maintenance, etc.

### GridTape enables rapid acquisition of an EM dataset of an adult *Drosophila* VNC

We acquired a dataset encompassing an adult female *Drosophila* VNC that consisted of 86.3 trillion voxels and spanned 21 million μm^3^ (Fig. 1D-J and Video S1). The dataset was captured from 4355 serial horizontal sections, each cut around 45 nm thick and collected onto GridTape continuously over 27 hours (22.1 seconds per section). Of these sections, 98% were positioned within 0.37 mm of the average section position, with only six sections having 20% or more of the VNC off the imageable slot area (Fig. 1E and S1F, see Methods). An additional three sections were lost before imaging due to support film breakage. No off-slot or lost sections were consecutive. Imaging required 60 continuous days on one TEMCA-GT at a rate of 42.73 ± 3.04 Mpixels per second (mean ± SD across sections) at 4.3 × 4.3 nm^2^ per pixel resolution. This amounted to 20.6 million images and 172.6 TB of raw 16-bit data. The dataset spans the VNC, from the subesophageal ganglion in the head through the neck connective to the thoracic ganglia where the leg, wing, haltere, and some neck motor neurons reside (Figs. 1D, F-H).

### Motor and sensory neurons occupy distinct domains within peripheral nerves

After aligning the acquired images into a three-dimensional image volume (see Methods), we searched for axon bundles leaving the VNC as peripheral nerves. We found all previously described nerves that innervate the legs, wings, halteres, and neck (Court et al., 2017; Power, 1948). For individual neurons passing through each nerve, we built skeleton models of their projection patterns within the VNC. Reconstructed neurons fell into three major morphological categories corresponding to motor, sensory, and central neurons (Baek and Mann, 2009; Brierley et al., 2012; Mamiya et al., 2018; Tsubouchi et al., 2017). Motor neurons had cell bodies located in the VNC, projected to a dorsal layer of the VNC, and did not contain synaptic vesicles or presynaptic specializations within the VNC neuropil (Fig. 2A and Video S2). Sensory neurons did not have cell bodies in the VNC, arborized more ventrally, and made synaptic outputs within the VNC neuropil (Fig. 2B and Video S2). Central neurons had a cell body in the VNC, made synaptic outputs in the VNC neuropil, and had projections confined to the central nervous system. An additional twenty neurons did not fall into one of the three main categories: the peripherally synapsing interneurons (King and Wyman, 1980), the octopaminergic dorsal unpaired median (DUM) neurons (Duch et al., 1999), and a new class of ‘multinerve’ neurons in the T1 neuromeres that project through multiple nerves (Fig. S2). Additionally, we counted 3738 axons travelling between the brain and VNC via the neck connective, consistent with previous counts (Coggshall et al., 1973).

**Figure 2.**
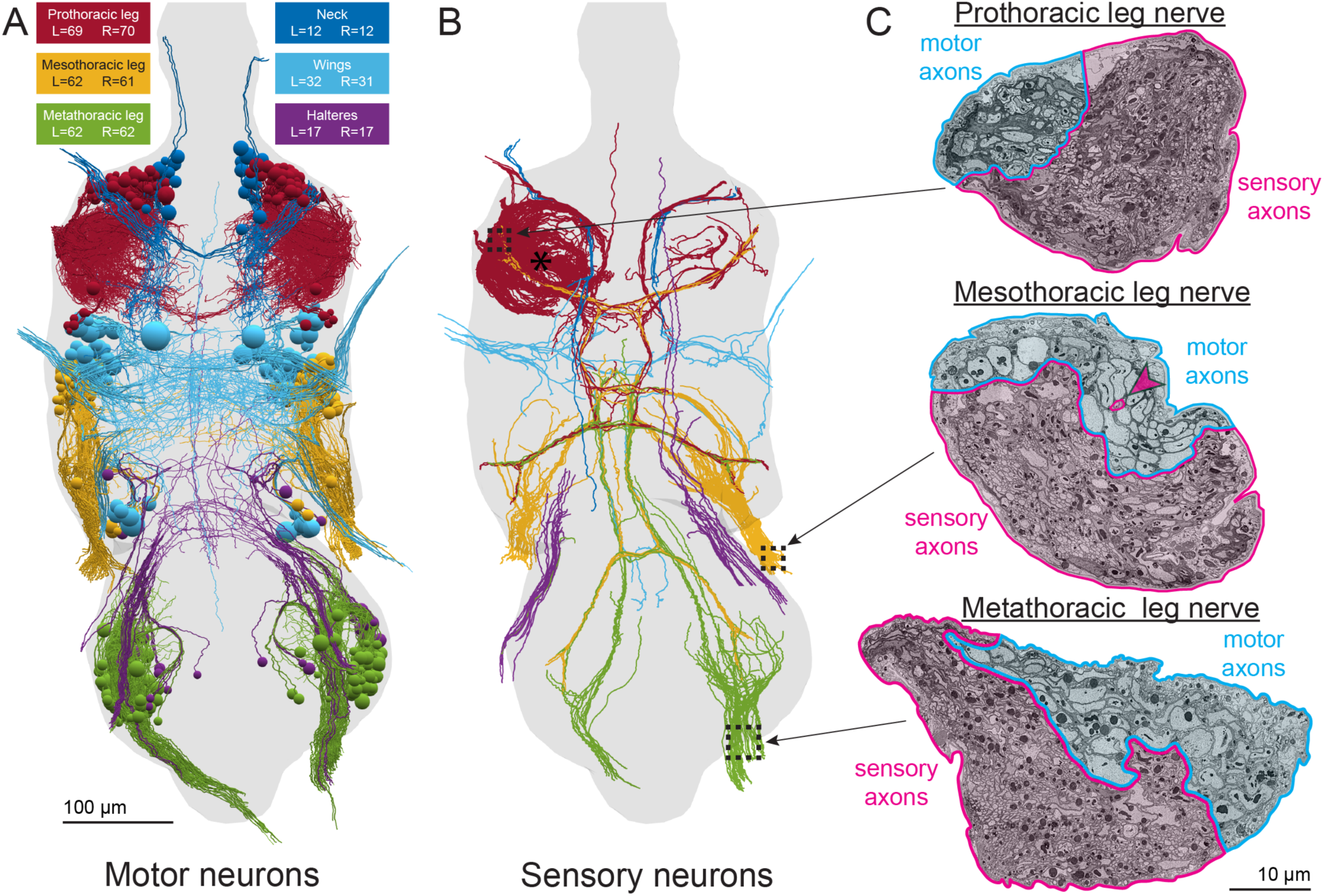
Reconstruction of motor and sensory neurons reveals precise functional domains in nerves. (A) All 507 motor neurons in the thoracic segments of the VNC were reconstructed from an EM dataset (Fig. 1). Each motor neuron projects its axon out one peripheral nerve, leaving the bounds of the EM dataset, to innervate muscles. Cell bodies are represented as spheres. An additional 34 motor neurons controlling neck muscles descend from the brain and could not be reconstructed back to their cell bodies (not shown). This rendering and all subsequent ones are viewed from the dorsal side of the VNC unless otherwise indicated. (B) Sensory axons entering the thoracic segments of the VNC. Sensory neurons have cell bodies outside the VNC, near their sensory organ. Reconstruction of sensory axons entering the VNC were focused primarily on the left T1 neuromere (asterisk). Same color code as (A). (C) Sections through the prothoracic, mesothoracic, and metathoracic leg nerves, which contain most of the sensory and motor axons connecting the VNC to the front, middle, and hind legs, respectively. Sections taken at the locations indicated by dashed boxes in (B). The leg nerves have distinct domains containing the axons of motor neurons (cyan) and the axons of sensory neurons (magenta). The only intermingling between motor and sensory axons is a group of three sensory axons within the motor domain of the mesothoracic leg nerve (magenta arrowhead). Scale bars, 100 µm (A-B), 10 µm (C).

We focused our reconstruction efforts on neurons projecting through the VNC’s peripheral nerves. We found that motor and sensory axons segregated into distinct spatial domains within leg nerves (Fig 2C) with only one exception, and consistent with findings in larger insects (Zill et al., 1980). Sensory axons outnumbered motor neuron axons by an order of magnitude in most nerves. For example, we found 867 sensory axons and 42 motor neuron axons in the left prothoracic leg nerve (ProLN) innervating the T1 neuromere. By reconstructing neurons that travel in the motor domain of each nerve, we identified a total of 507 motor neurons in the thoracic segments of the VNC. Together with 20 DUM neurons and two multinerve neurons (Fig S2), these reconstructions encompass the complete population of neurons that this VNC used to control the muscles of the legs, wings, halteres, and neck (Fig. 2A).

Due to the large number of sensory neurons entering the VNC, we next focused on reconstructing sensory neurons entering the left T1 neuromere. We reconstructed the main branches of 368 sensory neurons, focusing on proprioceptive sensory neurons. The sensory and motor neuron reconstructions presented here are made freely available, serving both as a database of cell types connecting the VNC to the body and as a starting point for future reconstruction efforts.

### Cell type-specific clustering of sensory and motor neuron axons

The reconstructed sensory and motor neurons fell into a number of morphological subtypes. Reconstructed sensory axons typically had one of four projection patterns, which corresponded to each of the four major sensory types (Tsubouchi et al., 2017; Tuthill and Wilson, 2016b): hair plate neurons (Merritt and Murphey, 1992), chordotonal neurons (Mamiya et al., 2018), bristle neurons (Murphey et al., 1989), and campaniform sensillum neurons (Merritt and Murphey, 1992) (Fig. 3 and Video S3, see Methods for classification criteria). We reconstructed the main branches of every proprioceptive axon in the left T1 neuromere, accounting for each major sensory neuron type other than bristle neurons. This included 33 hair plate neurons, 35 campaniform sensillum neurons, and 124 chordotonal neurons (Fig 3A and S3A). These counts are consistent with previous reports (Mamiya et al., 2018; Merritt and Murphey, 1992; Tsubouchi et al., 2017).

**Figure 3.**
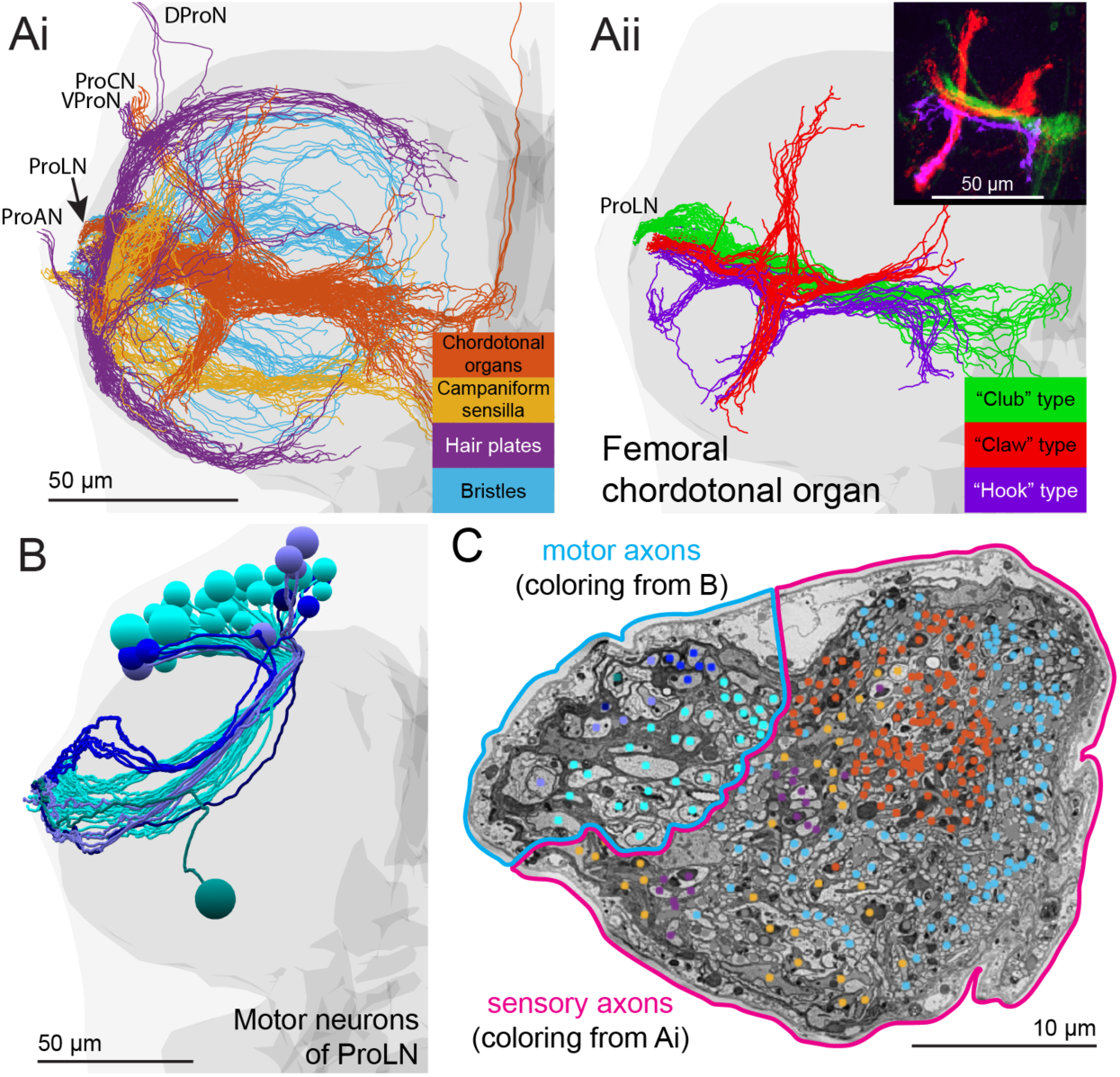
Subtypes of motor and sensory neurons are organized topographically in the VNC and peripheral nerves. (A) Reconstruction of the main branches of sensory neurons for the front left leg. (i) The four main functional subtypes of sensory neurons are identifiable based on their projection patterns in the VNC. The VNC is shaded light grey. The neuropil is shaded darker grey. The peripheral nerves are labeled: prothoracic accessory nerve (ProAN), prothoracic leg nerve (ProLN), ventral prothoracic nerve (VProN), prothoracic chordotonal nerve (ProCN), dorsal prothoracic nerve (DProN). (ii) Morphological subtypes of axons from the femoral chordotonal organ. Inset: These types were previously characterized using light microscopy. Different types encode different aspects of leg kinematics (adapted from (Mamiya et al., 2018)). (B) Reconstruction of the cell bodies and primary neurites of the 42 motor neurons of the left ProLN. Primary neurites travel through the neuromere in five distinct bundles (L1-L5, colored different shades of blue) before leaving the VNC. The five bundles contain 29, 6, 5, 1, and 1 neurons. See Figs. 4A and S3 for motor neuron bundles of other nerves. (C) Positions of the subtypes of sensory and motor neuron axons within the ProLN. Axons of each bundle from (B) are spatially clustered within the motor domain. Axons of sensory neurons also cluster by subtype within the nerve. Chordotonal axons, arising mostly from a single sensory organ in the leg (Mamiya et al. 2018), are more clustered than other types, which arise from sensory organs distributed across different segments of the leg (Tsubouchi et al., 2017). Scale bars, 50 µm (A-B), 10 µm (C).

The chordotonal neuron axons could be further subdivided into morphological subtypes that matched the “club”, “claw”, and “hook” neuron morphologies known to encode vibration, position, and direction of movement, respectively (Fig. 3Aii) (Mamiya et al., 2018). Five chordotonal axons also ascended directly to the brain (Tsubouchi et al., 2017).

By reconstructing motor neurons in the left T1 neuromere, we found that their primary neurites were spatially clustered, forming 18 distinct bundles (Fig. S3B). We hypothesize that these 18 bundles correspond to developmental lineages, which typically have their primary neurites bundled together (Lacin et al., 2019). Five of these bundles totaling 42 neurons exited the VNC through the ProLN (Fig. 3B), five bundles totaling 12 neurons through the prothoracic accessory nerve (ProAN), six bundles totaling 11 neurons through the ventral prothoracic nerve (VProN), and two bundles totaling four neurons through the dorsal prothoracic nerve (DProN; nomenclature from Court et al. 2017). We found a single bundle containing 29 motor neurons (cyan in Fig. 3B), likely corresponding to lineage 15B, reported to contain 28 motor neurons that mostly innervate muscles in the distal leg segments (Brierley et al., 2012). The remaining 17 clusters likely correspond to the remaining 14 motor neuron lineages (Baek and Mann, 2009).

In addition to having distinct morphologies within the T1 neuromere, subtypes of sensory and motor neuron axons were also topographically organized within the ProLN (Fig. 3C). Not only do motor and sensory axons occupy distinct regions of peripheral nerves (Fig. 2C), but this demonstrates a finer level of organization based on functional subtypes of sensory neurons and lineages of motor neurons.

### VNC atlas registration reveals uniqueness and symmetry of leg motor neurons

To examine the symmetry of motor neuron bundles, we next compared the populations of motor neurons controlling the left and right front legs. As observed for the 69 motor neurons for the front left leg, the 70 motor neurons for the right front leg formed 18 spatial clusters. These 18 bundles matched one-to-one between the left and right sides of the nervous system (Figs. 4A and S3B,D,E,F). The largest bundle on the right contained 30 neurons, one more than the 29 found in the largest bundle on the left, but the other 17 bundles contained identical numbers of motor neurons on the left and right sides.

**Figure 4.**
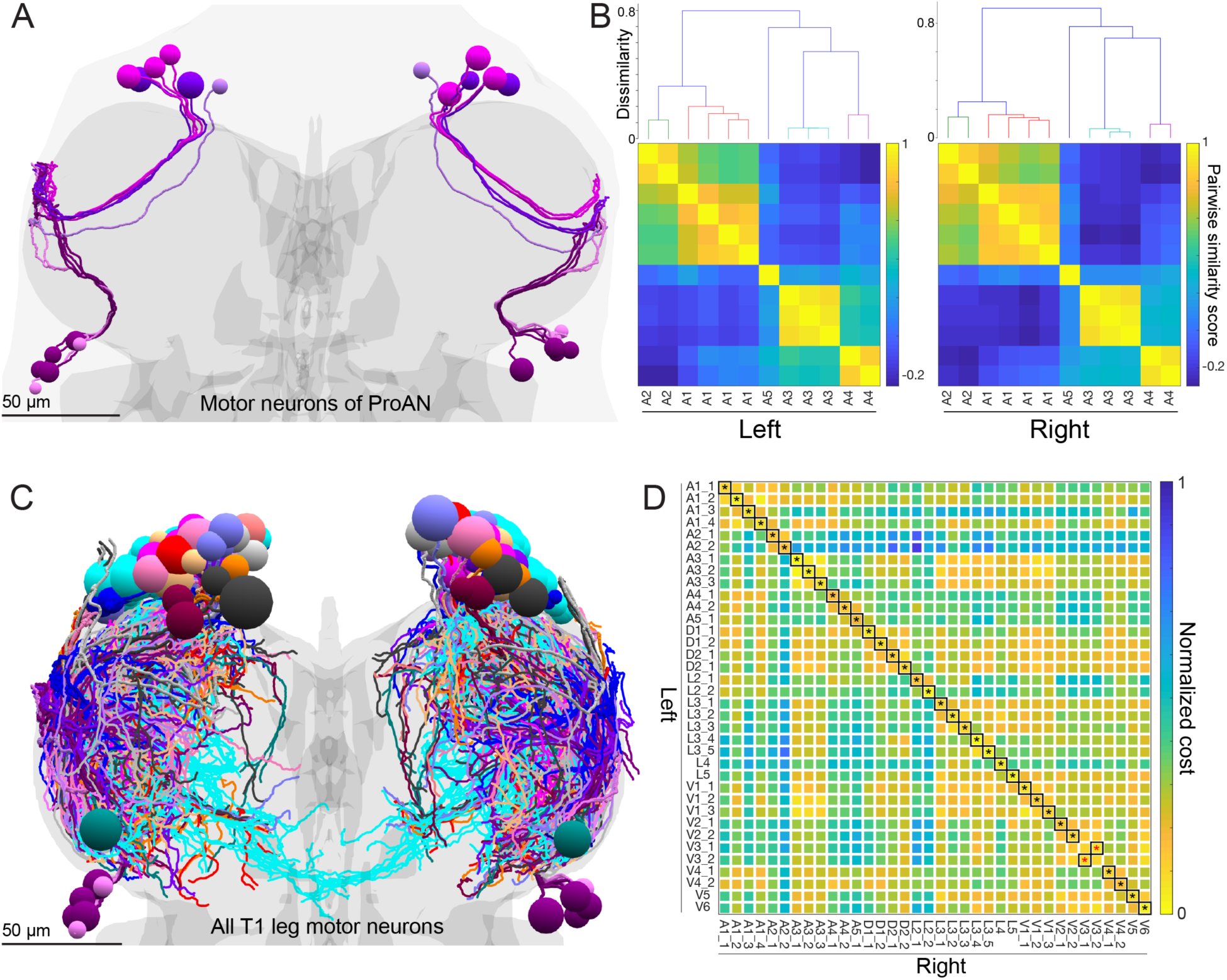
Motor neuron bundles, uniqueness, and symmetry. (A) Reconstruction of the cell bodies and primary neurites of the 24 motor neurons (12 per side) of the left and right ProAN. Primary neurites travel through the neuromere in five distinct and highly symmetric bundles (A1-5, colored in shades of purple). See also Fig. S3. (B) Quantitative analysis of motor neuron bundles. ProAN motor neurons on each side of the VNC were clustered based on the similarity of their primary neurites’ positions (NBLAST, see Methods). Top, dendrogram from hierarchical clustering. Members of each bundle from (A) cluster together. Bottom, matrix of pairwise similarity scores. Note the high degree of similarity between these quantitative representations of the motor neurons on the left and right side of the VNC. (C) The branching patterns of all 139 motor neurons arborizing in the T1 neuromeres were reconstructed. These reconstructions were then mapped into the VNC atlas coordinate system (see Fig. S4) to enable precise left-right comparisons across the midplane. (D) Identification of left-right homologous pairs of front leg motor neurons. Expert annotators identified 36 symmetric left-right pairs. A global pairwise assignment algorithm based on NBLAST similarity scores agreed (black asterisks) on all identified pairs except two (red asterisks). Scale bars, 50 µm (A, C).

To quantify this symmetry, we first registered the EM dataset to a standard VNC atlas to place the EM-reconstructed neurons into a reference coordinate system and to correct for asymmetries introduced by EM specimen preparation. The VNC atlas is a map of synapse density across the VNC (Bogovic et al., 2018). To register the EM dataset to the atlas, we first estimated the density of synapses across the EM dataset using an artificial neural network trained to identify synapses based on their ultrastructural features (Fig. S4, see Methods) (Buhmann et al., 2018). This allowed us to align the complete VNC EM volume to the adult female VNC atlas (Fig. S4D-E and Video S4) (Bogovic et al., 2018).

We were then able to quantitatively assess the similarity between neuronal morphologies using NBLAST (Costa et al., 2016) in the reference coordinate frame of the VNC atlas. As a proof-of-principle, we calculated similarity scores between the primary neurites of motor neurons exiting each nerve (Fig. S3C, see Methods) and performed hierarchical clustering on the scores. As expected, members of each of the 18 bundles clustered together on both the left and right sides (Fig. 4B and S3D-F). This confirmed that the motor neuron populations for the left and right front legs have their primary neurites organized systematically and symmetrically in bundles.

To investigate the ability to identify individual motor neurons and their left–right pairs, we reconstructed the branching patterns of all 139 motor neurons for the left and right front legs (Fig. 4C). Branches emerge from motor neurons’ primary neurites to form stereotyped arborizations. Motor neurons with particular branching patterns can often be recognized as innervating particular muscle groups (Baek and Mann, 2009; Brierley et al., 2012). We searched for bilaterally symmetric pairs of front leg motor neurons. We included motor neurons in bundles with equivalent numbers of neurons on the left and right sides. From this population of 40 neurons per side, we identified 36 left–right pairs by visual inspection, whose branching patterns appeared unique and symmetric (Video S5). To quantitatively verify these pairings, similarity scores were computed between left and right front leg motor neurons, after reflecting the latter across the midplane. From the similarity scores, we generated a globally optimal pairwise assignment (see Methods), which matched 34 of 36 (94%) of the predicted left–right pairs (Fig. 4D). In all cases, left–right pairs were members of symmetric primary neurite bundles. This result demonstrates that most motor neurons have an identifiable symmetric neuron on the opposite side of the VNC, with symmetric pairs likely controlling the same muscle in opposite legs (Baek and Mann, 2009). While left– right pairs were largely mirror symmetric, there was some variability in their branches. We often observed higher-order branches following slightly different paths to reach the same terminal zones (Fig S5A), consistent with findings in larval *Drosophila* (Schneider-Mizell et al., 2016).

These results illustrate the diversity of motor neurons and the symmetry of individual motor neuron morphologies. They also demonstrate the feasibility of identifying left–right copies of the same cell type within the VNC. Left–right comparisons can be helpful in revealing variations and compensations in developmental programs for specifying connectivity (Schneider-Mizell et al., 2016; Tobin et al., 2017).

### Bilaterally projecting leg sensory neurons co-activate motor neurons innervating different legs

Axons of campaniform sensillum neurons fell into three morphological categories, largely matching previously reported types in larger fly species (Merritt and Murphey, 1992). The first projects only to the neuromere for its leg of origin. The second projects to ipsilateral neuromeres corresponding to other legs on the same side of the body. The third category—which we call bilateral campaniform sensillum (bCS) neurons—project to multiple ipsilateral and contralateral neuromeres. bCS neurons had multiple striking features. First, this was the only type of leg sensory neuron to project across the midline (Fig. 5A-B). Second, bCS axons had the largest diameter of any leg sensory neuron, even exceeding the diameter of most motor neuron axons (Fig. 5C). By reconstructing the sensory neurons with the largest diameter axons for each leg, we identified 12 total bCS neurons in the VNC, with two originating in each of the six legs (Fig. 5A-B). All bCS neurons from the front legs projected to the front and middle leg neuromeres of the VNC (Fig 5Ai), those from the middle leg projected to all six neuromeres (Fig. 5Aii), and those from the hind legs projected to the hind and middle leg regions (Fig. 5Aiii). Notably, bCS axon branches were located directly alongside leg motor neuron axons (Fig. 5D). Therefore, we hypothesized that bCS neurons directly influence the activity of motor neurons across multiple legs in response to bCS stimulation of a single leg.

**Figure 5.**
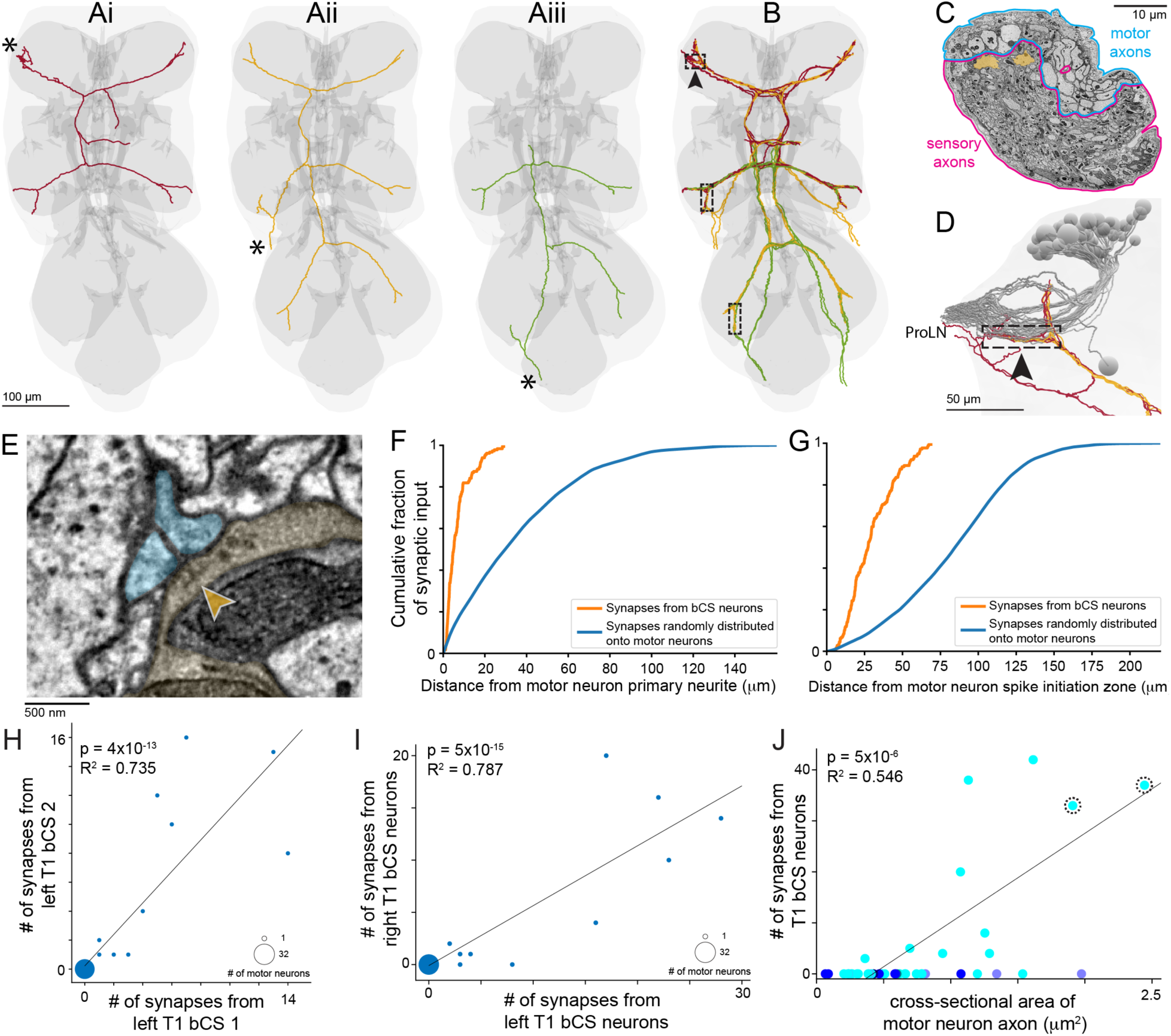
Bilateral campaniform sensillum (bCS) neurons directly connect to motor neurons with large-diameter axons near the spike-initiation zone. (A) Single bCS axons originating from the front (i), middle (ii), and hind (iii) left legs. Asterisks denote where each axon enters the VNC. (B) Two neurons with the morphologies show in (A) originate from each of the six legs, producing 12 total neurons of this type. bCS axons travel in the oblique tract (alongside other campaniform sensilla) in their neuromere of origin, and co-fasiculate with other bCS axons in a more dorsal tract in other neuromeres. Dashed boxes indicate a ∼50 µm-long tract in each neuromere where bCS axons originating from that neuromere travel alongside bCS axons originating from other neuromeres. (C) Right mesothoracic (T2) leg nerve. bCS axons (orange) are the largest-diameter leg sensory neuron, with diameters exceeding that of most leg motor neurons. The large diameter relative to motor neurons and other sensory neurons is observed in every leg nerve. (D) Lateral view of the location in left T1 indicated by the arrowhead in (B). Also shown are the cell bodies and primary neurites of motor neurons of the left ProLN (grey; same neurons as Fig. 3B). Note that in the boxed region, bCS axons originating from left and right T1 (red) and left and right T2 (yellow) all travel directly alongside primary neurites of ProLN motor neurons. (E) Synapse from a right T2 bCS axon (yellow) onto two left T1 motor neurons (cyan). The presynaptic T-bar structure (arrowhead), vesicles, and postsynaptic densities are visible. All 12 bCS neurons synapse onto motor neurons in each neuromere to which they project. (F-J) Analysis of all synaptic connections made by left and right T1 bCS axons along the 50 µm stretch shown in (D). (F) Distribution of distances from each bCS synapse to each postsynaptic motor neuron’s primary neurite (orange, n = 183 postsynaptic sites). bCS synapses directly target motor neuron primary neurites or short (<10 µm) branches coming off the primary neurite. Motor neuron arbors (Legend continued on next page) have many locations further from the primary neurite where input could be received but that the reconstructed bCS synapses do not target (blue, see Methods). (G) Distances from each bCS synapse to the putative motor neuron spike initiation zone (orange, n = 183 postsynaptic sites). bCS synapses densely innervate motor neurons at locations within 50 µm of the putative motor neuron spike initiation zone. Motor neuron arbors have many locations further from the spike initiation zone where input could be received but that the reconstructed bCS synapses do not target (blue, see Methods). (H) The two bCS axons originating from the left T1 leg (arbitrarily designated 1 and 2) target a subset of the left ProLN motor neurons. 32 out of 42 motor neurons receive no input from the reconstructed bCS axons. For any given motor neuron, the number of synaptic inputs from the two bCS axons are significantly correlated. (I) Connectivity from left T1 versus right T1 bCS axons onto left ProLN motor neurons. The two left bCS axons and two right bCS axons largely target the same motor neurons; note that the left and right T1 bCS axons most strongly target the same five motor neurons. (J) Relationship between the cross-sectional area of a motor neuron’s axon, the primary neurite bundle, and the number of synapses from bCS neurons. Points are colored according to which of the five ProLN bundles the motor neuron belongs to (same coloring as Fig. 3B). Only motor neurons in a single bundle receive any synapses from bCS neurons. Within that bundle, axon cross-sectional area is strongly correlated with number of synaptic inputs. Dashed circles indicate the two motor neurons with morphologies most similar to a functionally characterized fast motor neuron (see Fig. 6). Scale bars, 100 µm (A-B), 10 µm (C), 50 µm (D), 500 nm (E).

To investigate this hypothesis, we first asked whether bCS axons connect directly to leg motor neurons. Indeed, all 12 bCS axons made synapses directly onto motor neurons in each of the neuromeres to which they projected (Fig. 5E). To determine how frequently bCS synapses targeted motor neurons, we reconstructed all bCS synapses along the 50 µm-long stretch in left T1 where their axons travel alongside motor neuron primary neurites (Fig. 5D). In this region, left T1 bCS axons made 74 synapses and right T1 bCS axons made 43, of which 98.3% (115 of 117) had at least one motor neuron as a postsynaptic partner. There were 2.95 ± 1.15 (mean ± SD) postsynaptic partners at each synapse, totaling 345 postsynaptic sites. Of these, 60.3% were motor neurons, 23.8% were central neurons, and 15.9% could not be classified (see Methods). These connections were made either directly onto the motor neuron primary neurite or onto short (typically <10 µm) second-order branches (Fig. 5F). Furthermore, these connections were located within 50 µm of where each motor neuron’s primary neurite exits the VNC to become an axon (Fig. 5G). This location at the distal portion of the primary neurite is likely near the spike initiation zone, so bCS synapses appear well-positioned to drive activity in motor neurons (Gwilliam and Burrows, 1980).

These reconstructions revealed that the two bCS axons from the same leg synapse onto the same sub-population of ProLN motor neurons. Specifically, the number of synapses that a given ProLN motor neuron received from each of the two left T1 bCS neurons was highly correlated (Fig. 5H, R^2^ = 0.735, p = 4 × 10^−13^, *n* = 10 distinct motor neurons receiving 126 synapses). Inputs from the two right T1 bCS neurons were also correlated (R^2^ = 0.584, p = 4 × 10^−9^, *n* = 8 distinct motor neurons receiving 68 synapses). Notably, right and left T1 bCS axons synapsed onto the same motor neurons; the five motor neurons receiving the most synapses from left T1 bCS neurons were also the top five targets of the right T1 bCS neurons (Fig. 5I, R^2^ = 0.787, p = 7 × 10^−15^, *n* = 10 distinct motor neurons receiving 194 synapses).

The connections of bCS neurons are highly selective. They synapse onto only 10 left ProLN motor neurons, despite being positioned near most of the 42 ProLN motor neurons. What determines which motor neurons receive bCS input? First, bCS target only motor neurons within the largest bundle of 29 motor neurons (cyan in Fig. 3B), avoiding synapsing onto the 13 motor neurons in the other four bundles of ProLN motor neurons (Figs. 3-4). Moreover, within the targeted bundle, there was a strong correlation between the cross-sectional area of a motor neuron’s axon and the number of synapses it receives (Fig 5J, R^2^ = 0.546, p = 5 × 10^−6^, *n* = 29 motor neurons). Taken together, our results show that bCS axons from multiple legs converge to synapse onto a specific group of motor neurons with large-diameter axons, which are all members of the same bundle. Therefore, activity in a bCS neuron from either front leg likely stimulates synchronized, symmetric muscular contractions in both front legs.

### Fast flexor motor neurons are major postsynaptic targets of bCS neurons

Because bCS neurons synapse mainly onto motor neurons with large-diameter axons, we hypothesized that bCS neurons synaptically target “fast” motor neurons that control large ballistic movements. These fast motor neurons are distinct from the “slow” motor neurons that control small postural movements (Azevedo et al., 2019). To investigate this, we genetically targeted a fast motor neuron controlling the tibia flexor muscle of the front leg for whole-cell recording, filled it with dye via the whole-cell patch pipette, and imaged the cell and immunostained neuropil using confocal microscopy. We then traced the cell arbors in the VNC to produce a digital reconstruction from the light microscopy data. This reconstruction was then transformed into the same VNC atlas coordinate space to which the EM dataset was registered (Fig. S4). We repeated this process for a slow motor neuron also controlling a tibia flexor muscle of the front leg. This enabled us to quantitatively compare the morphological similarity between physiologically characterized neurons and EM-reconstructed neurons (Fig. 6).

**Figure 6.**
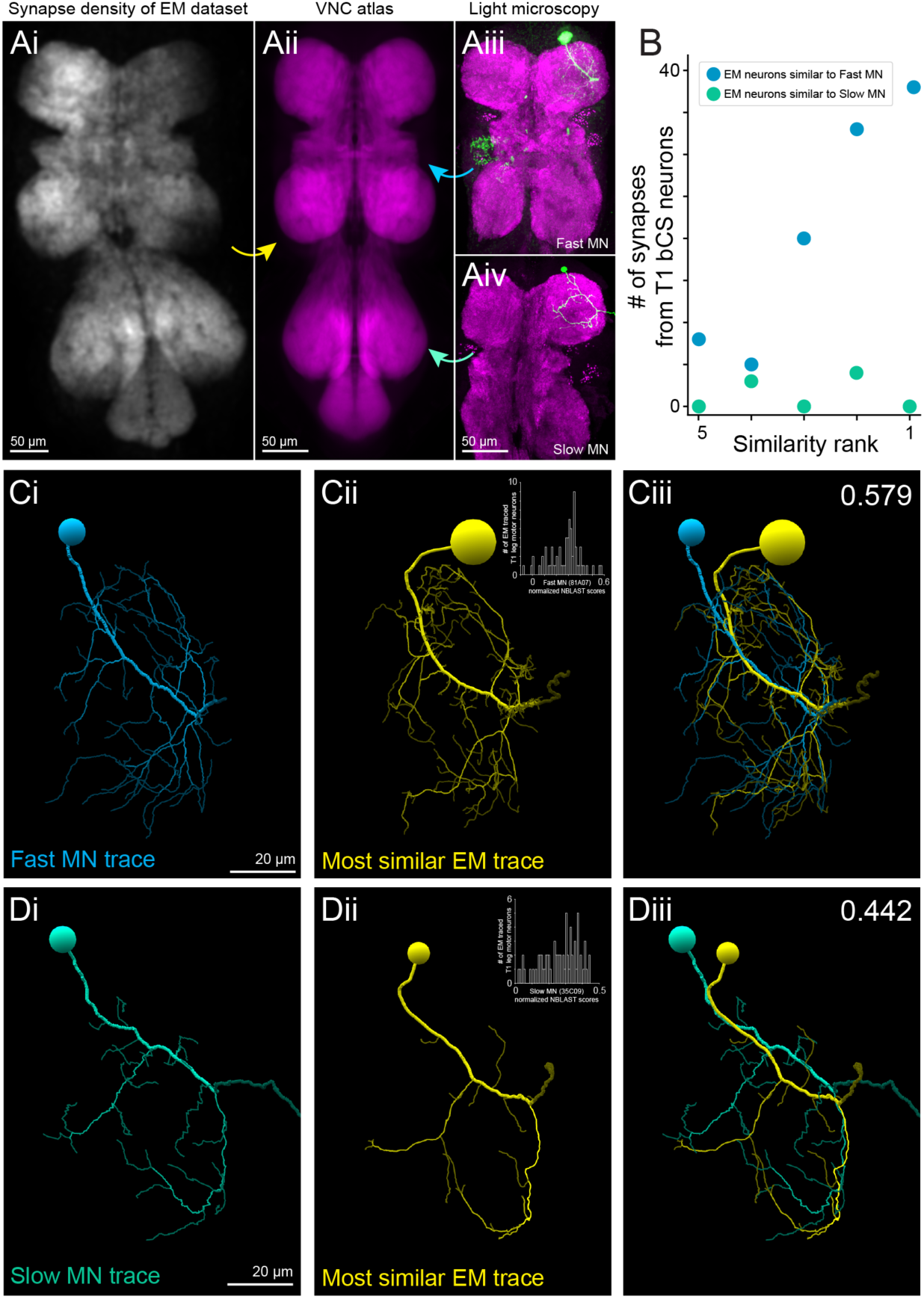
The fast tibia flexor motor neuron is a major synaptic target of bCS neurons. (A) Strategy for comparing morphologies of neurons imaged using light microscopy with neurons traced in EM. The synapse-density map of the EM dataset (Ai; see Fig. S4 for generation of this map) was registered to a standard VNC synapse-density map (Aii; Bogovic et al. 2018). Then, fluorescently labeled motor neurons (Aiii-iv) were also registered to the standard VNC for quantitative comparison with EM reconstructions using NBLAST (Costa et al. 2016). (B) Plot of the number of synapses from bCS neurons onto the top 5 most similar scoring EM-reconstructed neurons for the both fast (labeled by 81A07-Gal4) and slow (labeled by 35C09-Gal4) motor neurons. The EM neurons most similar to the fast motor neuron receive many bCS synapses, in contrast to the top matches for the slow motor neuron. See also Video S6. (C-D) Structure of light microscopy neurons and their most similar EM neurons. Displayed in the VNC atlas coordinate space with branches shaded by branching order (Strahler number). (Ci) Morphology of a fast tibia flexor motor neuron (labeled by 81A07-Gal4). The neuron was filled with dye following electrophysiological recording (Azevedo et al. 2019) and manually traced. (ii) EM-reconstructed motor neuron with the highest similarity score to the fast motor neuron. Inset: Histogram of NBLAST similarity scores between the fast motor neuron and the 69 EM-reconstructed motor neurons for the left front leg. (iii) Both (Ci) and (Cii) rendered together. NBLAST score of this pair in upper right corner. (D) Same as (C) for the slow motor neuron (green, labeled by 35C09-Gal4). Scale bars, 50 µm (A), 20 µm (C-D).

We used NBLAST (Costa et al., 2016) to search all 69 EM-reconstructed left front leg motor neurons for those with morphologies resembling the fast and slow tibia flexor motor neurons (Fig. 6C-D). For the fast motor neuron, we found that the highest-scoring motor neuron in the EM dataset had a highly similar structure (Fig. 6C). Moreover, the two top matching EM-reconstructed neurons both had axons with large cross-sectional areas and received many synaptic inputs (37 and 33 synapses) from T1 bCS neurons (Fig. 6B, dashed circles in Fig. 5J). All of these bCS inputs were near the putative spike initiation zones of the two motor neurons. For the slow motor neuron, the EM-reconstructed neurons with the highest similarity scores received very few synapses from bCS neurons (Fig. 6B,D and Video S6). Thus, we conclude that a major synaptic target of bCS neurons are fast motor neurons including a fast tibia flexor motor neuron.

These results illustrate the feasibility of automatically searching the VNC EM dataset for any reconstructed cell of interest. The starting point for this search is a confocal image of that cell and the surrounding neuropil. Because we registered the VNC EM dataset to a standard VNC atlas, this type of cross-modality search can be performed by any user.

## DISCUSSION

Large-scale neuronal wiring diagrams at single-synapse resolution will be a crucial element of future progress in neuroscience. Here, we present GridTape, a new technology for accelerating the acquisition of large-scale EM data. We demonstrated the power of this approach by acquiring a dataset encompassing an adult female *Drosophila* VNC. We then used this dataset to identify a novel monosynaptic circuit which directly links a specialized proprioceptor cell type with a specific set of motor neurons. We also illustrate a general pipeline for searching this dataset for cells of interest. The public release of this dataset represents a significant new resource, as well as an illustration of the capacity for rapid advances powered by this new technology.

### An accessible TEM pipeline for connectomics

Data acquisition has remained a rate-limiting step in the generation of large-scale EM datasets. Manual pickup of sections for TEM is slow, imprecise, and unreliable. Meanwhile, SEM-based approaches that circumvent the need for manual section collection have slow imaging speeds or require massive parallelization of expensive electron optics to acquire comparable datasets. The GridTape approach we developed increases the accessibility of EM connectomics for a wider scientific community. GridTape builds on previous efforts toward TEM parallelization and automation (Bock et al., 2011; Peltier et al., 2005; Zheng et al., 2018), but overcomes the need for manual pickup of sections for TEM, allowing much faster and more consistent section collection and imaging. Because sections are imaged nondestructively, GridTape is compatible with enhancement by post-section labeling and can benefit from multi-resolution re-imaging over large volumes (Hildebrand et al., 2017). By eliminating the need to separately handle thousands of fragile sections, GridTape reduces data loss and artifacts that compromise the quality of most TEM datasets. This results in better alignment of sections into a coherent, high signal-to-noise image volume, which leads to efficient and accurate reconstructions.

GridTape is also less expensive than high-throughput SEM platforms. For the current price of one commercial multi-beam SEM system (Eberle et al., 2015), ten TEMCA-GTs can be built, and samples collected on GridTape can be distributed across microscopes for simultaneous imaging. The fixed costs for the microscope hardware are accompanied by consumable costs associated with support film coating (currently ∼USD%4 per slot); however, we expect this cost to decrease due to further developments in tape coating technology and economies of scale.

In the future, GridTape acquisition rates will increase as cameras and imaging sensors continue to improve. Because TEM imaging is a widefield technique, imaging throughput can be increased by employing larger camera arrays and brighter electron sources. Moreover, sections larger than current slot dimensions (2 mm × 1.5 mm) could be accommodated by utilizing wider tape with larger slots, although custom microscopes may be necessary for very large samples and slot size will depend on material properties of the support film. However, GridTape should be compatible with thick sectioning approaches that subdivide tissue to overcome size limitations associated with other methods (Hayworth et al., 2015).

By enabling affordable, high-throughput EM imaging, GridTape makes it possible to study questions that require comparison of large-scale EM volumes from multiple individual organisms. These include questions related to development and ageing, sexual dimorphism, allelic variation, experience-dependent plasticity, the effects of perturbations, and disease mechanisms. This technology should also allow comparisons between the neuronal wiring diagrams of related species. By lowering the barriers to acquiring such datasets, this technology not only democratizes large-scale EM, but also changes the types of questions that it can be used to address.

### A complete adult *Drosophila* VNC dataset

In this study, we demonstrated the capabilities of the GridTape approach by generating an EM dataset of an adult female *Drosophila* VNC. This dataset provides a unique public resource for understanding how the *Drosophila* nervous system processes information to generate behavior. Additionally, it complements the recent release of an EM dataset comprising the complete adult female *Drosophila* brain (Zheng et al., 2018). Because the VNC comprises one third of the central nervous system, and because most brain functions are mediated via projections from the brain to the VNC (Namiki et al., 2018), it is critical to have the ability to study neuronal circuits in both the brain and the VNC at synaptic resolution. The dataset presented here includes all the intrinsic neurons of the VNC, all its sensory inputs, and all the descending and ascending axons connecting the brain to the VNC. We describe a straightforward pipeline for identifying any cell type of interest within the dataset by comparing reconstructions from the EM volume to confocal microscopy images (Fig. 6). Finally, as a foundation for future work in this dataset, we make publicly available a large set of annotated sensory and motor neuron reconstructions (Figs. 2-4).

### Direct sensory feedback to motor neurons

Because EM datasets include all the cells within a volume of interest, they permit the discovery of novel cell types and synaptic connections that may be overlooked by other methods. By performing a targeted reconstruction of sensory afferents, we found that the leg sensory neurons with the largest-diameter axons are the bilaterally projecting campaniform sensilla, which make direct synapses onto leg motor neurons (Fig. 5). These direct connections are specific to the motor neuron with the largest-caliber axons including a fast tibia flexor neuron (Figs. 5J, 6), which is involved in fast, ballistic movements. This is evidence that speed is essential for the function of these connections. These synapses are specifically located near the putative motor neuron spike initiation zone (Fig. 5G), suggesting bCS inputs are poised to influence motor neuron spiking.

The unique bilateral and intersegmental projections of bCS neurons suggests that they are capable of directly influencing multiple limbs on both sides of the body (Fig. 5A-B). This leads to several hypotheses about their function. Prior work has suggested that campaniform sensilla encode information about step timing that could be used to drive the transition between stance and swing phases of walking (Dallmann et al., 2017; Ridgel et al., 1999). However, we observe that bCS neurons synapse onto the same motor neurons on both sides of the body (Fig 5I), indicating that they likely support symmetric movements of the legs. This makes it less likely that the bCS neurons contribute to anti-phasic walking movements. Instead, bCS neurons seem well-positioned to underlie a fast reflex where multiple legs flex in response to bCS activation. Campaniform sensilla can signal either increases or decreases in load, depending on the sensillum’s placement and orientation on the leg (Zill and Moran, 1981; Zill et al., 1981). Therefore, bCS neuron activation could serve to forcefully stabilize posture in response to additional weight (e.g. to prevent the body from being crushed) or to grip a surface in response to a loss of load on the legs (e.g. to prevent being blown away by a gust of wind). Whether bCS neurons signal increases or decreases in load will require future functional studies.

Monosynaptic sensory-to-motor neuron connectivity is infrequent in the larval *Drosophila* nervous system (Zarin et al., 2019), but more frequent in other adult insects (Burrows, 1996). One possibility is that direct sensory feedback in adults is key for control of segmented limbs, enabling precise and adaptive limb movements. Such connections being absent in larvae may indicate that controlling a limbless body relies less on sensory feedback and more on feedforward processing. Another possibility is that adult movements simply occur on faster timescales than do larval movements, so having fast monosynaptic sensory feedback is particularly useful when the musculature is able to bring about fast movements. Indeed, research on escape responses has demonstrated that high-velocity movements are often brought about by the fastest neuronal pathways (Eaton et al., 1977; Trimarchi and Schneiderman, 1995).

### Diversity and stereotypy within complete leg motor neuron populations

Motor neurons have diverse but stereotyped functions, reflecting the array of muscles and muscle fibers they innervate. Some motor neurons have unique and reproducible transcription factor signatures that underlie their physiological properties and axonal morphology (Enriquez et al., 2015; Venkatasubramanian et al., 2019). These unique transcription factor patterns specify morphologies that are fairly stereotyped across animals (Baek and Mann, 2009; Brierley et al., 2012).

Our results extend these previous studies to show that many leg motor neurons are sufficiently stereotyped that they are individually identifiable by structure alone, and that homologous pairs of motor neurons on opposite sides of the body are readily matched. Because we reconstructed a complete population of leg motor neurons in our EM dataset, we were able to show that mirror symmetry is a systematic and general principle that applies to this cell type (Fig. 4). In contrast, neurons at the sensory periphery seem to have more redundant copies and variable copy numbers (Takemura et al., 2015; Tobin et al., 2017).

Although motor neurons are sufficiently stereotyped to be identifiable as left–right pairs, individual branches of these neurons can reach their terminal zones via variable routes (Fig. S5). This type of branch-specific variation has been described previously in *Drosophila* larvae (Schneider-Mizell et al., 2016). These variations suggest limits to the precision of the molecular and genetic programs that guide dendritic and axonal branching during development.

### Adult *Drosophila* as a model system for studying circuit mechanisms of motor control

Detailed, comprehensive connectivity patterns within the nerve cord were previously acquired in organisms without limbs. These include *C. elegans* (White et al., 1986), leeches (Stent et al., 1978), lampreys (Buchanan and Grillner, 1987; Grillner, 2003), and *Drosophila* larvae (Cardona et al., 2010; Fushiki et al., 2016; Ohyama et al., 2015; Zwart et al., 2016). These studies have enabled a greater mechanistic understanding of how the nervous system controls locomotor rhythms that produce swimming and crawling.

However, less is known about neuronal connectivity underlying motor control in limbed species. Coordinated limb movement for locomotion and posture requires motor control and coordination at multiple levels, from coordinating antagonist muscles of individual joints to coordinating multiple joints in a given limb to coordinating multiple limbs for effective locomotor behavior. Here, we present a connectomic dataset that will enable complete mapping of connectivity of the neuronal circuits that control the legs and wings of an adult *Drosophila melanogaster*. Combined with recent advances in the ability to record activity from genetically identified VNC neurons during behavior (Azevedo et al., 2019; Chen et al., 2018; Mamiya et al., 2018), we expect that a greater understanding of the circuit basis for complex motor control is within reach.

## ACKNOWLEDGEMENTS

We thank A. DeCoursey, M. Liu, T. Pedersen, R. Xu, A. Yeager for neuron tracing; the HMS EM Core for technical support; S. Gerhard, G. Hood, X. Chen and R. Zheng for programming and software support; A. Bleckert, D. Brittain, R. Torres, N. da Costa, and R.C. Reid for their feedback and support; K. Hayworth for his vision, inspiration, and advice; J. Lichtman, R. Schalek, and K. Hayworth for an ATUM device and SEM imaging support; O. Mazor and P. Gorelik for engineering support; C. Harvey, L. Cheadle, and M. Greenberg for tissue samples; R. Fetter for histology advice; J. Bogovic for VNC atlas alignment advice; N. Perrimon, L. Ventakasubramanian, R. Mann, S. Rayshubskiy and R. Wilson for sharing *Drosophila* lines; H. Somhegyi for assistance with dissections; M. Pecot for providing fly husbandry resources; T. Ayers, R. Smith, and Luxel Corporation for coating tape; and R. Wilson, E. Raviola, and the Lee laboratory for comments on the manuscript. This work was supported by NIH grants (R21NS085320, RF1MH114047, RF1MH117808), the Bertarelli Program in Translational Neuroscience and Neuroengineering, Edward R. and Anne G. Lefler Center, Stanley and Theodora Feldberg Fund, a Genise Goldenson Award, and the Intelligence Advanced Research Projects Activity (IARPA) of the Department of Interior/Interior Business Center (DoI/IBC) through contract number D16PC00004 to W-C.A.L.; and by NIH grants (T32MH20017, T32HL007901) for D.G.C.H. Portions of this research were conducted on the Orchestra High Performance Compute Cluster at Harvard Medical School partially provided through NIH NCRR grant 1S10RR028832-01. The views and conclusions contained herein are those of the authors and should not be interpreted as representing the official policies or endorsements, either expressed or implied, of the funding sources including NIH, IARPA, DoI/IBC, or the U.S. Government.

## AUTHOR CONTRIBUTIONS

J.M-S., J.C.T., and W-C.A.L. conceptualized the biological project. D.G.C.H. and W-C.A.L. conceptualized the technology. B.J.G. and D.G.C.H. designed, built, and developed software for tape milling. B.J.G, D.G.C.H and B.L.S. designed, built, and developed software for the ATUM and ultramicrotome modifications. B.J.G. designed, built, and developed software for tape handling; computerized microscope control; and the reel-to-reel GridTape imaging stage. B.L.S. developed software for tape staining and the reel-to-reel GridTape imaging stage. B.J.G., D.G.C.H., B.L.S., J.M-S., L.A.T., and A.T.K. developed instrumentation control and analysis software. J.M-S. and W-C.A.L. prepared samples. D.G.C.H. and J.M-S. sectioned samples. J.M-S. collected EM data. A.W.A. and J.C.T. collected cell fills and confocal microscopy data. J.B. and J.F. developed the synaptic partner prediction network. T.N., L.A.T., and J.B. applied the synaptic partner prediction network to predict presynaptic sites in the EM dataset. J.M-S. stitched and aligned the EM dataset and aligned the light microscopy and EM dataset to the reference atlas. J.M-S. and W-C.A.L. performed analysis of the reconstruction data. J.M-S., D.G.C.H., B.J.G., and W-C.A.L. wrote the paper with input from other authors.

## METHODS

### Animals and tissue preparation

All procedures involving animals were conducted in accordance with the ethical guidelines of the NIH and approved by the IACUC at Harvard Medical School. The Standing Committee on the Use of Animals in Research and Training of Harvard University approved all animal experiments.

We fixed and stained the central nervous system of one adult female *Drosophila melanogaster* (aged 1-2 days post-eclosion, genotype y,w/w[1118]; +; P{VT025718-Gal4}attP2/P{pBI-UASC–3×MYC– sbAPEX2–dlg-S97}18). Following fixation (2% paraformaldehyde/2.5% glutaraldehyde) and dissection (Tobin et al., 2017), the specimen was reacted with diaminobenzadine (DAB) and H_2_O_2_ as described previously (Zhang et al., 2019), but an EM-dense label was not observed in this sample. The dissected central nervous system was then post-fixed and stained with 1% osmium tetroxide/1.5% potassium ferrocyanide, followed by 1% thiocarbohydrazide, a subsequent incubation in 2% osmium tetroxide, then 1% uranyl acetate, followed by lead aspartate (Walton, 1979), then dehydrated with a graded ethanol series. The specimen was then embedded in epoxy resin (TAAB 812 Epon, Canemco), positioned in a cutout of mouse cortex processed for EM using the same protocol without the DAB reaction. The mouse thalamus specimen (Fig. S1H) was prepared as previously described (Deerinck et al., 2010; Hua et al., 2015) and post-section stained with stabilized lead citrate (Ultrastain II, Leica). The VNC sections were not post-section stained following sectioning onto GridTape.

*Drosophila* Gal4 lines and husbandry for matching physiologically characterized cells in the EM dataset are described in (Azevedo et al., 2019). Genotypes for the flies used for searching against EM traced cells (Fig. 6) were: w[1118]; P{JFRC7-20XUAS-IVS-mCD8::GFP} attp40/+; P{y[+t7.7] w[+mC]=GMR81A07-GAL4}attP2/+ and w[1118]; P{JFRC7-20XUAS-IVS-mCD8::GFP} attp40/+; P{y[+t7.7] w[+mC]=GMR35C09-GAL4}attP2/+.

### Substrate production

GridTape was produced from 125 µm-thick aluminum-coated Kapton® film (Dunmore) slit into 8 mm-wide reels of 35 m length (Metlon Corporation). This stock tape was modified using a custom laser-milling system consisting of a reel-to-reel tape positioning machine and commercial 1 W ultraviolet laser marking system (Samurai, DPSS Lasers). Control software triggered laser milling of a 30 mm length of tape, used custom computer vision to check the result of the cutting, advanced the tape 30 mm and finally adjusted the position of the tape to align the next 30 mm of tape to cut. This system enabled the autonomous production of long, >30 m, lengths of cut tape containing over 5000 slots. Following laser milling, the cut tape was cleaned by wiping it with isopropyl alcohol-soaked lint free wipes. Finally, the cut tape was coated with a 50 nm-thick TEM support film (LUXFilm®, Luxel Corporation). A description of the GridTape system was previously posted as a preprint (Graham et al., 2019).

### Sample block trimming

In preparation for sectioning, embedded tissue blocks were trimmed (Trim 90, Diatome) into an oblong hexagonal shape (Fig. 1C) with 3.5–4 mm height, 1–2 mm width, a greater than 90° degree bottom angle and less than 90° top angle.

### Serial sectioning

An ultramicrotome (UC7, Leica) and diamond knife (4 mm, 35° Ultra or Ultra-Maxi, Diatome) were used to cut ultra-thin serial sections (∼45 nm) from prepared samples. These sections were collected using a modified automated tape-collecting ultramicrotome (ATUM; (Hayworth et al., 2014)). All tape guides and rollers on the ATUM were modified by adding a 4 mm-wide trough to prevent contact with the TEM support film spanning GridTape slots. Additionally, an optical interrupter (GP1A57HRJ00F, Sharp Electronics) was affixed to the ATUM to detect the passage of GridTape slots (Fig. S1C), and a hall-effect sensor (A1301EUA-T, Allegro MicroSystems) and magnet were attached to the microtome swing arm to detect the cutting of sections (Fig. S1B). Custom software monitored the period and relative phase-offset of these two sensors during section collection. By setting the microtome to a fixed cutting speed and varying the ATUM tape speed, effective phase-locking at a fixed offset was achieved (Fig. S1D). For this specimen to reach stable sectioning conditions, an initial stretch of 45 sections was collected while adjustments were made to the tape speed and fixed offset. Of these 45 sections, 21 were off-slot and thus not imageable with TEM. The 24 that were on-slot contained small portions of the abdominal ganglion and were imaged and included in the dataset. Of the 4355 serial sections subsequently collected, the VNC region was completely off-slot in two sections and partially off-slot in four sections (20%, 30%, 70%, and 90% off-slot). Due to support film breakage, three sections were completely lost before imaging, and four were partially lost (10%, 10%, 20%, and 40%). One additional section was partially lost (10%) because it cut very thinly and a portion was distorted. No further sections had substantial data loss. Note that sections collected onto GridTape but off-slot can still be acquired using the traditional ATUM-SEM approach (Fig. S1H). Because of the reliability of the section placement (Fig. S1F), SEM imaging was not required for the VNC dataset.

### Measuring section placement consistency

Section placement was measured by first capturing photographs (Flea3 FL3-U3-13E4C-C, PointGrey) of each slot. Collimated low-angle illumination (MWWHL4, Thorlabs) enhanced the visibility of sections adhered to the tape. Using the captured images, the location of the slot was first found using the Fiji plugin “Template Matching and Slice Alignment” (https://sites.google.com/site/qingzongtseng/template-matching-ij-plugin), selecting the slot as the template. Any failures to automatically find the slot (<1% occurrence) were corrected manually in Fiji (Schindelin et al., 2012). Subsequently, the location of the tissue section was found using the same plugin, selecting a prominent feature of the tissue section as the template. The VNC tissue’s shape and appearance changed significantly across the 4355 section series, so template matching was performed on smaller batches of ∼500–1000 sections, with a separate feature chosen for template matching in each batch. This approach enabled automatic identification of the tissue’s placement for ∼98% of sections. The remaining ∼2% of sections that were not correctly identified were located manually in Fiji. The sections needing this manual correction mainly fell into two categories: sections that were cut extremely thinly, causing the tissue to have reduced visibility, or sections with the template feature placed near the slot edge.

### TEM imaging

To perform TEM imaging of sections collected onto GridTape, a custom in-vacuum, reel-to-reel stage was constructed (Fig. S1G) and attached to a TEMCA (Bock et al., 2011) consisting of a TEM (JEOL 1200 EX) with a 2×2 array of sCMOS cameras (Zyla 4.2, Andor). The stage allows a 7500-slot, 45 m-long roll of GridTape to be loaded into the microscope for imaging under vacuum. After loading and pump-down, a set of pinch drives (one on each side of the TEM column) allows linear movement of GridTape to exchange and position sections under the electron beam in preparation for imaging. After positioning, both pinch drives dispense a small amount of GridTape towards the center of the column, introducing slack on both sides of the sample held under the beam. This allows an XY stack of piezo nanopositioners (SLC-1720, SmarAct) to make the many small movements necessary to montage large areas. At 4.3 nm lateral resolution, the TEMCA field of view for a single location was just over 16µm square. By capturing many images at slightly overlapping regions (typically 20–30%) for a single section, millimeter-sized regions of interest could be imaged. Imaging regions for each section were selected using the photographs described in the section above using a custom graphical user interface in MATLAB (MathWorks). Magnification at the microscope was 2500×, accelerating potential was 120 kV, and beam current was ∼90 µA through a tungsten filament. The VNC dataset was acquired at a net sustained imaging rate of 42.73 ± 3.04 Mpixels per second (mean ± SD across sections), equivalent to a “burst” imaging rate of ∼160 Mpixels per second for a single microscope.

### Section stitching and serial-section alignment

Image alignment was performed with a custom pipeline that deployed AlignTK’s image alignment functions (https://mmbios.pitt.edu/aligntk-home) in parallel on a computing cluster (Bock et al., 2011; Lee et al., 2016; Tobin et al., 2017). After acquisition, camera images for each section were virtually stitched together into seamless montages. All section-to-section alignment was performed on 8x downscaled versions of these section montages. To align the 4355 stitched sections into a three-dimensional volume, an initial volume was first generated comprised of every 25^th^ section. The only features recognizable across gaps of 25 sections were neuronal nuclei, so this initial volume positioned every 25^th^ section in a location that ensured a given nucleus would stay at the same (x,y) location across the ∼150 sections in which each nucleus was visible. This positioning of every 25^th^ section was used as a global constraint on the full dataset’s alignment (absolute_maps option in AlignTK’s align function).

Due to the small number of sections with artifacts or missing data, elastic alignment (AlignTK’s register function) between neighboring sections was sufficient for generating a high-quality global alignment, except for 27 sections where alignment to direct neighbors and second neighbors was necessary. Additionally, no sections were mis-ordered, eliminating the need for a section order correction step. Overall, the consistency of GridTape section collection simplified the alignment process substantially and enabled the final volume to have high quality alignment (Video S1).

### Neuron reconstruction

We reconstructed neurons in the EM dataset as described previously (Lee et al., 2016; Tobin et al., 2017). We used CATMAID (Saalfeld et al., 2009) to manually place a series of marker points down the middle of each neurite to generate skeletonized models of neuronal arbors. We annotated neurons passing through each peripheral nerve and reconstructed those neurons into the VNC. Neurons that had a cell body in the VNC and arborized in the neuropil were considered motor neurons. Neurons that made synaptic outputs in the neuropil but did not have cell bodies in the VNC were considered sensory neurons. Neurons with projections and cell bodies in the VNC but that did not pass through a peripheral nerve were considered central neurons. We identified synapses using a combination of ultrastructural criteria, specifically the existence of a presynaptic T-bar, presynaptic vesicles, and postsynaptic densities. This procedure follows a previously described and validated protocol for reconstructing neurons in serial section TEM datasets (Schneider-Mizell et al., 2016).

We were able to identify cell types for VNC sensory and motor neurons by their stereotyped projection patterns, which corresponded well with previous observations of these neurons using light microscopy (Baek and Mann, 2009; Brierley et al., 2012; Mamiya et al., 2018; Merritt and Murphey, 1992; Tsubouchi et al., 2017). Bristle neuron axons traveled along either the anterior, posterior, or ventral edge of the neuromere without significant branching. Hair plate neuron axons bifurcated or trifurcated upon entering the VNC and projected along the anterior, posterior, and lateral edges of the neuromere. Chordotonal neuron axons projected through the middle of the neuromere toward the midline. Campaniform sensillum axons projected down the oblique tract, located posterior to the chordotonal neuron axons.

In peripheral nerves, axons of motor neurons were clustered together. After finding a single motor axon in a given nerve, we reconstructed its neighbors, continuing to reconstruct further neighbors until all motor neurons in the nerve were reconstructed. We confirmed that sensory neurons near the motor domain were in fact sensory neurons by reconstructing them into the VNC, and we additionally reconstructed large-caliber axons in the sensory domain that we suspected could be motor neurons despite their position. For the left prothoracic leg nerve, we confirmed 324 of the 867 axons in the sensory domain were sensory neurons according to the criteria above (Figs. 2C, 3A,C). No motor neuron axons have yet been found in the sensory domain of any peripheral nerve. We found one case where three sensory neurons had axons located in the motor domain of the right mesothoracic leg nerve (Figs. 2C). We used this reconstruction approach to identify all motor neurons in all thoracic nerves (i.e. all nerves except the abdominal nerves), and we reconstructed all 507 thoracic motor neurons from their primary neurites back to their cell bodies. We then reconstructed most of the microtubule-containing backbones (Schneider-Mizell et al., 2016) of all front leg motor neurons. Reconstructions proceeded until multiple expert annotators were able to independently identify left-right homologous pairs of front leg motor neurons by their symmetrical morphology. Annotators were blind to the left-right pair predictions generated by analysis of NBLAST similarity scores (Fig. 4D).

### Automated synapse prediction and VNC atlas alignment

To transform EM reconstructions into the atlas space, we computationally generated a “neuropil stain” (Heinrich et al., 2018) by automatically detecting presynaptic specializations in the EM volume that would be labeled by immunostaining (Kittel et al., 2006). Specifically, we trained and deployed a convolutional neural network to automatically identify presynaptic locations across the entire EM dataset (Buhmann et al., 2018). We trained a neural network on section-wise, four-fold downsampled raw data (effective voxel size 17.2×17.2×45 nm). We used a 3D U-Net (Falk et al., 2019), comprised of four resolution levels with downsample factors in *x, y, z* of (2, 2, 1), (2, 2, 1), and (2, 2, 3). The topmost level contained four feature maps and the number of feature maps in subsequent levels increases with a factor of five. Convolutional passes were comprised of two convolutions with kernel sizes of (3, 3, 3) followed by a rectified linear unit (ReLU) activation. A final convolution with kernel size (1, 1, 1) produced the map of predicted presynaptic sites. We trained the network on randomly chosen crop-outs of size (172, 172, 42) from the CREMI challenge training data (https://cremi.org) and additional annotations from the VNC dataset (three densely annotated ground-truth cubes of 3×3×3 µm (768×768×75 voxels) and 13 ground-truth cubes with no synapses), which we augmented with random *x,y*-transpositions, *x,y*-flips, continuous rotations around the *z*-axis, and section-wise elastic deformations and intensity changes. We used mean-squared loss to train the network on ground-truth presynaptic specialization maps, which we generated by rasterizing the presynaptic point annotations as spheres with a radius of 100 nm.

We chose a standard VNC atlas (Bogovic et al., 2018) as our reference coordinate space. We used the presynaptic specializations predicted in the EM volume to register the EM dataset to the atlas. The density of the predicted synapses matched the spatial extents of the VNC neuropil. Regions of low synaptic density corresponded to fasciculated neuronal tracts devoid of synapses (Fig. S4B-C). We subsequently downsampled and blurred (σ = 2 µm) the predicted synapse locations in EM to match the resolution of the light microscopy data (Fig. S4D), then aligned this data to the atlas using elastix (http://elastix.isi.uu.nl/) (Video S4). Confocal microscopy data was also transformed into the same VNC atlas coordinate system using elastix (Fig. 6).

### Clustering and symmetry analysis

For the primary neurite clustering analysis (Figs. 4B and S3D-F), EM-reconstructed neurons were first transformed into the VNC atlas space using the registration described above. Then, neurons were pruned to exclude any parts of the reconstruction falling outside the VNC neuropil. This retained neurons’ neurites in the neuropil, but excluded their cell bodies, which are known to have variable positions from fly to fly even for identified neurons and are therefore not reliable indicators of a neuron’s identity (Baek and Mann, 2009). Neurons were further pruned to only include their primary neurite (see Fig S3C). NBLAST similarity scores (Costa et al., 2016) were calculated between each pair of pruned neurons in both forward and reverse directions (i.e. neuron A to neuron B and neuron B to neuron A) and normalized such that the similarity score of each neuron with itself is equal to 1. The forward and reverse scores were then averaged to generate a final similarity score for each pair of neurons. Hierarchical clustering with single linkage was performed on similarity scores for motor neurons of each peripheral nerve using the SciPy Python package. The clustering dendrograms and neuron reconstructions were visually inspected, and a cut height on each dendrogram was chosen that separated motor neuron bundles traveling along distinct trajectories.

For symmetry analysis (Fig. 4D), neurons were also transformed into the VNC atlas space and pruned to exclude their cell bodies as described above, but all branches emerging from the primary neurite were included instead of being pruned. Neurons on the right side of the dataset were reflected across the midplane of the atlas to enable comparison with neurons on the left side. NBLAST similarity scores were calculated as described above between each left-side motor neuron and each reflected right-side motor neuron. Because higher NBLAST scores reflect greater morphological similarity, to compute a cost metric for dissimilarity, we subtracted each pairwise score from the maximum score so that the most similar (highest scoring) pair had zero cost. Based on these costs, we used the Munkres algorithm in MATLAB (MathWorks) to compute a globally optimal pairwise assignment between individual motor neurons on the left and right sides of the VNC (Munkres, 1957).

### Analysis of synaptic connectivity for bCS neurons

All output synapses were identified in the two bCS axons arising from the left prothoracic leg nerve and in the two arising from the right prothoracic leg nerve along the ∼50µm branch in the left T1 neuromere indicated in Figure 5D. Multiple independent annotators reviewed these synapse identifications to ensure accuracy and completeness. Postsynaptic partners of each synapse were reconstructed until each one could be identified as a motor neuron (by connecting to an existing motor neuron reconstruction), a central neuron (by making a synaptic output), or until reconstruction could not be continued due to uncertainty in where the neuron continued. We never observed a sensory neuron being postsynaptic to a bCS synapse. The postsynaptic motor neurons included 10 ProLN motor neurons in the L1 bundle, and one VProN motor neuron. Analysis in Figures 5F-G was restricted to the seven ProLN motor neurons receiving five or more synapses from bCS neurons. Analysis in Figures 5H-J included all ProLN motor neurons.

Analysis was carried out in Python using pymaid (https://pymaid.readthedocs.io/en/latest/index.html) for pulling reconstructions from CATMAID, SciPy for linear regression, and matplotlib for visualization. For measuring distances between synapses and particular locations on motor neurons (Fig. 5F-G), geodesic or “along-the-arbor” distance was always calculated. To determine the distribution of distances between possible synaptic locations and the primary neurite, we computed the distances from all positions on the motor neuron arbor to the primary neurite (Fig. 5F), and we assumed that all locations on the motor neuron were equally likely to receive synaptic input. In reality, synapses are preferentially positioned on the distal branches of neurons (Schneider-Mizell et al., 2016), so the random distributions presented here likely underestimate the distances from the primary neurite at which synaptic inputs are found. This means the bias of bCS synapses to target regions close to the primary neurite relative to randomly positioned input is likely even stronger than suggested by this analysis.

For calculating distances to the putative spike initiation zone, the distal-most branch point of each motor neuron’s primary neurite was used as the approximate location of the spike initiation zone. Primary neurites branch extensively as they travel through the VNC neuropil, but then stop branching as they leave the VNC to become an axon. The spike initiation zone is located near this transition point between primary neurite and axon (Gwilliam and Burrows, 1980). To determine a distribution of distances between possible synaptic locations and the spike initiation zone, we computed the distances from all positions on the motor neuron arbor to this distal-most branch point (Fig. 5G).

To measure the cross-sectional area of left ProLN motor neuron axons (Fig. 5J), we selected five sections spanning a 600 section (27 µm) range where the ProLN traveled directly perpendicular to the sectioning plane. In each of the five sections, the polygon selection tool in Fiji was used to measure the area of each of the 42 ProLN motor neuron axons. Measured areas were averaged across the five sections.

Linear regression (Figs. 5H-J), including the determination of the line of best fit, R^2^ value, and p-value, was performed using scipy.stats.linregress. For Figures 5H-I, all ProLN motor neurons were included in the regression. For Figure 5J, only the motor neurons in the largest ProLN bundle (cyan) were included in the regression.

### Light microscopy-based cell matching

Intracellular labeling, immunohistochemistry, confocal microscopy, and tracing of genetically identified cells (Fig. 6) was performed as described in (Azevedo et al., 2019). Briefly, targeted neurons were labeled during whole-cell patch pipette recordings with 13 mM neurobiotin in the internal solution. After whole-cell recordings, the dissected VNC was lightly fixed in 4% paraformaldehyde in phosphate-buffered saline (PBS) for 20 min. The tissue was then washed in PBST (PBS + Triton, 0.2% w/w), incubated in blocking solution (PBST + 5% normal goat serum) for 20 min, and then incubated for 24 hrs in blocking solution containing a primary antibody for neuropil counterstain (1:50 mouse anti-Bruchpilot, Developmental Studies Hybridoma Bank, nc82). After a subsequent PBST wash, the tissue was incubated in blocking solution containing secondary antibodies for 24 hr (streptavidin AlexaFluor conjugate, Invitrogen; 1:250 goat anti-mouse AlexaFluor conjugate, Invitrogen). Following staining, the tissue was mounted in Vectashield (Vector Labs) and confocal stacks were acquired using a Zeiss 510 confocal microscope. Cells were traced in Fiji (Schindelin et al., 2012), using the Simple Neurite Tracing plugin (Longair et al., 2011). Neuron traces were transformed into the VNC atlas space using elastix.

### Data and code availability

The EM dataset and reconstructions will be available at: https://vnc1.catmaid.virtualflybrain.org/. Reel-to-reel instrumentation designs and software will be available at: https://www.lee.hms.harvard.edu/resources or https://github.com/htem/GridTapeStage. Additional code is available at https://github.com/htem or upon reasonable request.

## REFERENCES

Azevedo, A.W., Dickinson, E.S., Gurung, P., Venkatasubramanian, L., Mann, R., and Tuthill, J.C. (2019). A size principle for leg motor control in *Drosophila*. bioRxiv, 730218.

Baek, M., and Mann, R.S. (2009). Lineage and birth date specify motor neuron targeting and dendritic architecture in adult *Drosophila*. J Neurosci 29, 6904–6916.

Bock, D.D., Lee, W.C.A., Kerlin, A.M., Andermann, M.L., Hood, G., Wetzel, A.W., Yurgenson, S., Soucy, E.R., Kim, H.S., and Reid, R.C. (2011). Network anatomy and *in vivo* physiology of visual cortical neurons. Nature 471, 177–182.

Bogovic, J.A., Otsuna, H., Heinrich, L., Ito, M., Jeter, J., Meissner, G., Nern, A., Colonell, J., Malkesman, O., Ito, K., et al. (2018). An unbiased template of the *Drosophila* brain and ventral nerve cord. bioRxiv, 376384.

Brierley, D.J., Rathore, K., VijayRaghavan, K., and Williams, D.W. (2012). Developmental origins and architecture of *Drosophila* leg motoneurons. The Journal of Comparative Neurology 520, 1629–1649.

Briggman, K.L., Helmstaedter, M., and Denk, W. (2011). Wiring specificity in the direction-selectivity circuit of the retina. Nature 471, 183–188.

Buchanan, J.T., and Grillner, S. (1987). Newly identified ‘glutamate interneurons’ and their role in locomotion in the lamprey spinal cord. Science 236, 312–314.

Buhmann, J., Krause, R., Lentini, R.C., Eckstein, N., Cook, M., Turaga, S., and Funke, J. (2018). Synaptic partner prediction from point annotations in insect brains. Paper presented at: International Conference on Medical Image Computing and Computer-Assisted Intervention (Springer).

Burrows, M. (1996). The neurobiology of an insect brain (Oxford University Press on Demand).

Buschges, A., Akay, T., Gabriel, J.P., and Schmidt, J. (2008). Organizing network action for locomotion: insights from studying insect walking. Brain Res Rev 57, 162–171.

Cardona, A., Saalfeld, S., Preibisch, S., Schmid, B., Cheng, A., Pulokas, J., Tomancak, P., and Hartenstein, V. (2010). An integrated micro- and macroarchitectural analysis of the *Drosophila* brain by computer-assisted serial section electron microscopy. PLoS Biol 8.

Chen, C.L., Hermans, L., Viswanathan, M.C., Fortun, D., Aymanns, F., Unser, M., Cammarato, A., Dickinson, M.H., and Ramdya, P. (2018). Imaging neural activity in the ventral nerve cord of behaving adult *Drosophila*. Nat Commun 9, 4390.

Coggshall, J., Boschek, C., and Buchner, S. (1973). Preliminary Investigations on a Pair of Giant Fibers in the Central Nervous System of Dipteran Flies. Zeitschrift für Naturforschung C 28, 783–784b.

Costa, M., Manton, J.D., Ostrovsky, A.D., Prohaska, S., and Jefferis, G.S. (2016). NBLAST: Rapid, Sensitive Comparison of Neuronal Structure and Construction of Neuron Family Databases. Neuron 91, 293–311.

Court, R.C., Armstrong, J.D., Borner, J., Card, G., Costa, M., Dickinson, M., Duch, C., Korff, W., Mann, R., Merritt, D., et al. (2017). A Systematic Nomenclature for the *Drosophila* Ventral Nervous System. bioRxiv.

Dallmann, C.J., Hoinville, T., Durr, V., and Schmitz, J. (2017). A load-based mechanism for inter-leg coordination in insects. Proc Biol Sci 284.

Deerinck, T.J., Bushong, E.A., Thor, A., and Ellisman, M.H. (2010). NCMIR methods for 3D EM: a new protocol for preparation of biological specimens for serial block face scanning electron microscopy. Microscopy, 6–8.

Dickerson, B.H., de Souza, A.M., Huda, A., and Dickinson, M.H. (2019). Flies Regulate Wing Motion via Active Control of a Dual-Function Gyroscope. Curr Biol 29, 3517–3524 e3513.

Duch, C., Mentel, T., and Pfluger, H.J. (1999). Distribution and activation of different types of octopaminergic DUM neurons in the locust. The Journal of Comparative Neurology 403, 119–134.

Eaton, R.C., Bombardieri, R.A., and Meyer, D.L. (1977). The Mauthner-initiated startle response in teleost fish. J Exp Biol 66, 65–81.

Eberle, A.L., Mikula, S., Schalek, R., Lichtman, J., Tate, M.L.K., and Zeidler, D. (2015). High-resolution, high-throughput imaging with a multibeam scanning electron microscope. J Microsc 259, 114–120.

Enriquez, J., Venkatasubramanian, L., Baek, M., Peterson, M., Aghayeva, U., and Mann, R.S. (2015). Specification of individual adult motor neuron morphologies by combinatorial transcription factor codes. Neuron 86, 955–970.

Falk, T., Mai, D., Bensch, R., Cicek, O., Abdulkadir, A., Marrakchi, Y., Bohm, A., Deubner, J., Jackel, Z., Seiwald, K., et al. (2019). U-Net: deep learning for cell counting, detection, and morphometry. Nat Methods 16, 67–70.

Fushiki, A., Zwart, M.F., Kohsaka, H., Fetter, R.D., Cardona, A., and Nose, A. (2016). A circuit mechanism for the propagation of waves of muscle contraction in *Drosophila*. Elife 5.

Graham, B.J., Hildebrand, D.G.C., Kuan, A.T., Maniates-Selvin, J.T., Thomas, L.A., Shanny, B.L., and Lee, W.-C.A. (2019). High-throughput transmission electron microscopy with automated serial sectioning. bioRxiv, 657346.

Grillner, S. (2003). The motor infrastructure: from ion channels to neuronal networks. Nat Rev Neurosci 4, 573–586.

Gwilliam, G., and Burrows, M. (1980). Electrical characteristics of the membrane of an identified insect motor neurone. Journal of Experimental Biology 86, 49–61.

Hayworth, K.J., Morgan, J.L., Schalek, R., Berger, D.R., Hildebrand, D.G., and Lichtman, J.W. (2014). Imaging ATUM ultrathin section libraries with WaferMapper: a multi-scale approach to EM reconstruction of neural circuits. Front Neural Circuits 8, 68.

Hayworth, K.J., Xu, C.S., Lu, Z., Knott, G.W., Fetter, R.D., Tapia, J.C., Lichtman, J.W., and Hess, H.F. (2015). Ultrastructurally smooth thick partitioning and volume stitching for large-scale connectomics. Nat Methods 12, 319–322.

Heinrich, L., Funke, J., Pape, C., Nunez-Iglesias, J., and Saalfeld, S. (2018). Synaptic Cleft Segmentation in Non-isotropic Volume Electron Microscopy of the Complete Drosophila Brain (Cham: Springer International Publishing).

Hildebrand, D.G.C., Cicconet, M., Torres, R.M., Choi, W., Quan, T.M., Moon, J., Wetzel, A.W., Scott Champion, A., Graham, B.J., Randlett, O., et al. (2017). Whole-brain serial-section electron microscopy in larval zebrafish. Nature 545, 345–349.

Hua, Y., Laserstein, P., and Helmstaedter, M. (2015). Large-volume en-bloc staining for electron microscopy-based connectomics. Nat Commun 6, 7923.

Kanning, K.C., Kaplan, A., and Henderson, C.E. (2010). Motor neuron diversity in development and disease. Annu Rev Neurosci 33, 409–440.

Kasthuri, N., Hayworth, K.J., Berger, D.R., Schalek, R.L., Conchello, J.A., Knowles-Barley, S., Lee, D., Vazquez-Reina, A., Kaynig, V., Jones, T.R., et al. (2015). Saturated Reconstruction of a Volume of Neocortex. Cell 162, 648–661.

Kiehn, O. (2011). Development and functional organization of spinal locomotor circuits. Curr Opin Neurobiol 21, 100–109.

King, D.G., and Wyman, R.J. (1980). Anatomy of the giant fibre pathway in Drosophila. I. Three thoracic components of the pathway. J Neurocytol 9, 753–770.

Kittel, R.J., Wichmann, C., Rasse, T.M., Fouquet, W., Schmidt, M., Schmid, A., Wagh, D.A., Pawlu, C., Kellner, R.R., Willig, K.I., et al. (2006). Bruchpilot promotes active zone assembly, Ca2+ channel clustering, and vesicle release. Science 312, 1051–1054.

Knott, G., Marchman, H., Wall, D., and Lich, B. (2008). Serial section scanning electron microscopy of adult brain tissue using focused ion beam milling. J Neurosci 28, 2959–2964.

Kornfeld, J., Benezra, S.E., Narayanan, R.T., Svara, F., Egger, R., Oberlaender, M., Denk, W., and Long, M.A. (2017). EM connectomics reveals axonal target variation in a sequence-generating network. Elife 6.

Lacin, H., Chen, H.M., Long, X., Singer, R.H., Lee, T., and Truman, J.W. (2019). Neurotransmitter identity is acquired in a lineage-restricted manner in the *Drosophila* CNS. Elife 8.

Lee, T.J., Kumar, A., Balwani, A.H., Brittain, D., Kinn, S., Tovey, C.A., Dyer, E.L., da Costa, N.M., Reid, R.C., Forest, C.R., et al. (2018). Large-scale neuroanatomy using LASSO: Loop-based Automated Serial Sectioning Operation. PLoS One 13, e0206172.

Lee, W.C.A., Bonin, V., Reed, M., Graham, B.J., Hood, G., Glattfelder, K., and Reid, R.C. (2016). Anatomy and function of an excitatory network in the visual cortex. Nature 532, 370–374.

Longair, M.H., Baker, D.A., and Armstrong, J.D. (2011). Simple Neurite Tracer: open source software for reconstruction, visualization and analysis of neuronal processes. Bioinformatics 27, 2453–2454.

Mamiya, A., Gurung, P., and Tuthill, J.C. (2018). Neural Coding of Leg Proprioception in *Drosophila*. Neuron.

Merk, A., Bartesaghi, A., Banerjee, S., Falconieri, V., Rao, P., Davis, M.I., Pragani, R., Boxer, M.B., Earl, L.A., Milne, J.L.S., et al. (2016). Breaking Cryo-EM Resolution Barriers to Facilitate Drug Discovery. Cell 165, 1698–1707.

Merritt, D.J., and Murphey, R.K. (1992). Projections of leg proprioceptors within the CNS of the fly *Phormia* in relation to the generalized insect ganglion. The Journal of Comparative Neurology 322, 16–34.

Miller, A. (1950). The internal anatomy and histology of the imago of *Drosophila melanogaster*. In The Biology of Drosophila, M. Demerec, ed. (New York, NY: John Wiley & Sons), pp. 420–531.

Morgan, J.L., Berger, D.R., Wetzel, A.W., and Lichtman, J.W. (2016). The Fuzzy Logic of Network Connectivity in Mouse Visual Thalamus. Cell 165, 192–206.

Munkres, J. (1957). Algorithms for the assignment and transportation problems. Journal of the society for industrial and applied mathematics 5, 32–38.

Murphey, R.K., Possidente, D., Pollack, G., and Merritt, D.J. (1989). Modality-specific axonal projections in the CNS of the flies *Phormia* and *Drosophila*. The Journal of Comparative Neurology 290, 185–200.

Namiki, S., Dickinson, M.H., Wong, A.M., Korff, W., and Card, G.M. (2018). The functional organization of descending sensory-motor pathways in *Drosophila*. Elife 7.

Niven, J.E., Graham, C.M., and Burrows, M. (2008). Diversity and evolution of the insect ventral nerve cord. Annu Rev Entomol 53, 253–271.

O’Sullivan, A., Lindsay, T., Prudnikova, A., Erdi, B., Dickinson, M., and von Philipsborn, A.C. (2018). Multifunctional Wing Motor Control of Song and Flight. Curr Biol 28, 2705–2717 e2704.

Ohyama, T., Schneider-Mizell, C.M., Fetter, R.D., Aleman, J.V., Franconville, R., Rivera-Alba, M., Mensh, B.D., Branson, K.M., Simpson, J.H., Truman, J.W., et al. (2015). A multilevel multimodal circuit enhances action selection in *Drosophila*. Nature 520, 633–639.

Peltier, S., Bouwer, J., Jin, L., Khodjasaryan, K., Geist, S., Xuong, N., and Ellisman, M. (2005). Design of a new 8k x 8k lens coupled detector for wide-field, high-resolution transmission electron microscopy. Microsc Microanal, 610–611.

Power, M.E. (1948). The thoracico-abdominal nervous system of an adult insect, Drosophila melanogaster. The Journal of Comparative Neurology 88, 347–409.

Pringle, J. (1938). Proprioception in insects: II. The action of the campaniform sensilla on the legs. Journal of Experimental Biology 15, 114–131.

Ridgel, A.L., Frazier, S.F., Dicaprio, R.A., and Zill, S.N. (1999). Active signaling of leg loading and unloading in the cockroach. Journal of Neurophysiology 81, 1432–1437.

Saalfeld, S., Cardona, A., Hartenstein, V., and Tomancak, P. (2009). CATMAID: collaborative annotation toolkit for massive amounts of image data. Bioinformatics 25, 1984–1986.

Schindelin, J., Arganda-Carreras, I., Frise, E., Kaynig, V., Longair, M., Pietzsch, T., Preibisch, S., Rueden, C., Saalfeld, S., Schmid, B., et al. (2012). Fiji: an open-source platform for biological-image analysis. Nat Methods 9, 676–682.

Schmidt, H., Gour, A., Straehle, J., Boergens, K.M., Brecht, M., and Helmstaedter, M. (2017). Axonal synapse sorting in medial entorhinal cortex. Nature 549, 469–475.

Schneider-Mizell, C.M., Gerhard, S., Longair, M., Kazimiers, T., Li, F., Zwart, M.F., Champion, A., Midgley, F.M., Fetter, R.D., Saalfeld, S., et al. (2016). Quantitative neuroanatomy for connectomics in *Drosophila*. Elife 5.

Sjostrand, F.S. (1958). Ultrastructure of retinal rod synapses of the guinea pig eye as revealed by three-dimensional reconstructions from serial sections. J Ultrastruct Res 2, 122–170.

Soler, C., Daczewska, M., Da Ponte, J.P., Dastugue, B., and Jagla, K. (2004). Coordinated development of muscles and tendons of the *Drosophila* leg. Development 131, 6041–6051.

Stent, G.S., Kristan, W.B., Jr., Friesen, W.O., Ort, C.A., Poon, M., and Calabrese, R.L. (1978). Neuronal generation of the leech swimming movement. Science 200, 1348–1357.

Strausfeld, N.J., Seyan, H., and Milde, J. (1987). The neck motor system of the fly *Calliphora erythrocephala*-I. Muscles and motor neurons. Journal of Comparative Physiology A 160, 205–224.

Takemura, S.Y., Bharioke, A., Lu, Z., Nern, A., Vitaladevuni, S., Rivlin, P.K., Katz, W.T., Olbris, D.J., Plaza, S.M., Winston, P., et al. (2013). A visual motion detection circuit suggested by *Drosophila* connectomics. Nature 500, 175–181.

Takemura, S.Y., Xu, C.S., Lu, Z., Rivlin, P.K., Parag, T., Olbris, D.J., Plaza, S., Zhao, T., Katz, W.T., Umayam, L., et al. (2015). Synaptic circuits and their variations within different columns in the visual system of *Drosophila*. Proc Natl Acad Sci U S A 112, 13711–13716.

Tapia, J.C., Wylie, J.D., Kasthuri, N., Hayworth, K.J., Schalek, R., Berger, D.R., Guatimosim, C., Seung, H.S., and Lichtman, J.W. (2012). Pervasive synaptic branch removal in the mammalian neuromuscular system at birth. Neuron 74, 816–829.

Tobin, W.F., Wilson, R.I., and Lee, W.C.A. (2017). Wiring variations that enable and constrain neural computation in a sensory microcircuit. Elife 6.

Trimarchi, J.R., and Schneiderman, A.M. (1995). Flight initiations in *Drosophila melanogaster* are mediated by several distinct motor patterns. J Comp Physiol A 176, 355–364.

Tsubouchi, A., Yano, T., Yokoyama, T.K., Murtin, C., Otsuna, H., and Ito, K. (2017). Topological and modality-specific representation of somatosensory information in the fly brain. Science 358, 615–623.

Tuthill, J.C., and Azim, E. (2018). Proprioception. Curr Biol 28, R194–R203.

Tuthill, J.C., and Wilson, R.I. (2016a). Mechanosensation and adaptive motor control in insects. Current Biology.

Tuthill, J.C., and Wilson, R.I. (2016b). Parallel transformation of tactile signals in central circuits of *Drosophila*. Cell 164, 1046–1059.

Venkatasubramanian, L., Guo, Z., Xu, S., Tan, L., Xiao, Q., Nagarkar-Jaiswal, S., and Mann, R.S. (2019). Stereotyped terminal axon branching of leg motor neurons mediated by IgSF proteins DIP-alpha and Dpr10. Elife 8.

Walton, J. (1979). Lead aspartate, an en bloc contrast stain particularly useful for ultrastructural enzymology. J Histochem Cytochem 27, 1337–1342.

Wanner, A.A., Genoud, C., Masudi, T., Siksou, L., and Friedrich, R.W. (2016). Dense EM-based reconstruction of the interglomerular projectome in the zebrafish olfactory bulb. Nat Neurosci 19, 816–825.

White, J.G., Southgate, E., Thomson, J.N., and Brenner, S. (1986). The structure of the nervous system of the nematode *Caenorhabditis elegans*. Phil Trans Royal Soc London B 314, 1–340.

Xu, C.S., Hayworth, K.J., Lu, Z., Grob, P., Hassan, A.M., Garcia-Cerdan, J.G., Niyogi, K.K., Nogales, E., Weinberg, R.J., and Hess, H.F. (2017). Enhanced FIB-SEM systems for large-volume 3D imaging. Elife 6.

Zarin, A.A., Mark, B., Cardona, A., Litwin-Kumar, A., and Doe, C.Q. (2019). A *Drosophila* larval premotor/motor neuron connectome generating two behaviors via distinct spatio-temporal muscle activity. bioRxiv, 617977.

Zhang, Q., Lee, W.C.A., Paul, D.L., and Ginty, D.D. (2019). Multiplexed peroxidase-based electron microscopy labeling enables simultaneous visualization of multiple cell types. Nat Neurosci 22, 828–839.

Zheng, Z., Lauritzen, J.S., Perlman, E., Robinson, C.G., Nichols, M., Milkie, D., Torrens, O., Price, J., Fisher, C.B., Sharifi, N., et al. (2018). A Complete Electron Microscopy Volume of the Brain of Adult *Drosophila melanogaster*. Cell 174, 730–743 e722.

Zill, S.N., and Moran, D.T. (1981). The exoskeleton and insect proprioception. I. Responses of tibial campaniform sensilla to external and muscle-generated forces in the American cockroach, *Periplaneta americana*. Journal of Experimental Biology 91, 1–24.

Zill, S.N., Moran, D.T., and Varela, F.G. (1981). The exoskeleton and insect proprioception: II. Reflex effects of tibial campaniform sensilla in the American cockroach, *Periplaneta americana*. Journal of Experimental Biology 94, 43–55.

Zill, S.N., Underwood, M.A., Rowley, J.C., 3rd, and Moran, D.T. (1980). A somatotopic organization of groups of afferents in insect peripheral nerves. Brain Res 198, 253–269.

Zwart, M.F., Pulver, S.R., Truman, J.W., Fushiki, A., Fetter, R.D., Cardona, A., and Landgraf, M. (2016). Selective Inhibition Mediates the Sequential Recruitment of Motor Pools. Neuron 91, 615–628.

