## Supplementary Materials for "Reconstruction of motor control circuits in adult *Drosophila* using automated transmission electron microscopy"

### SUPPLEMENTAL FIGURES

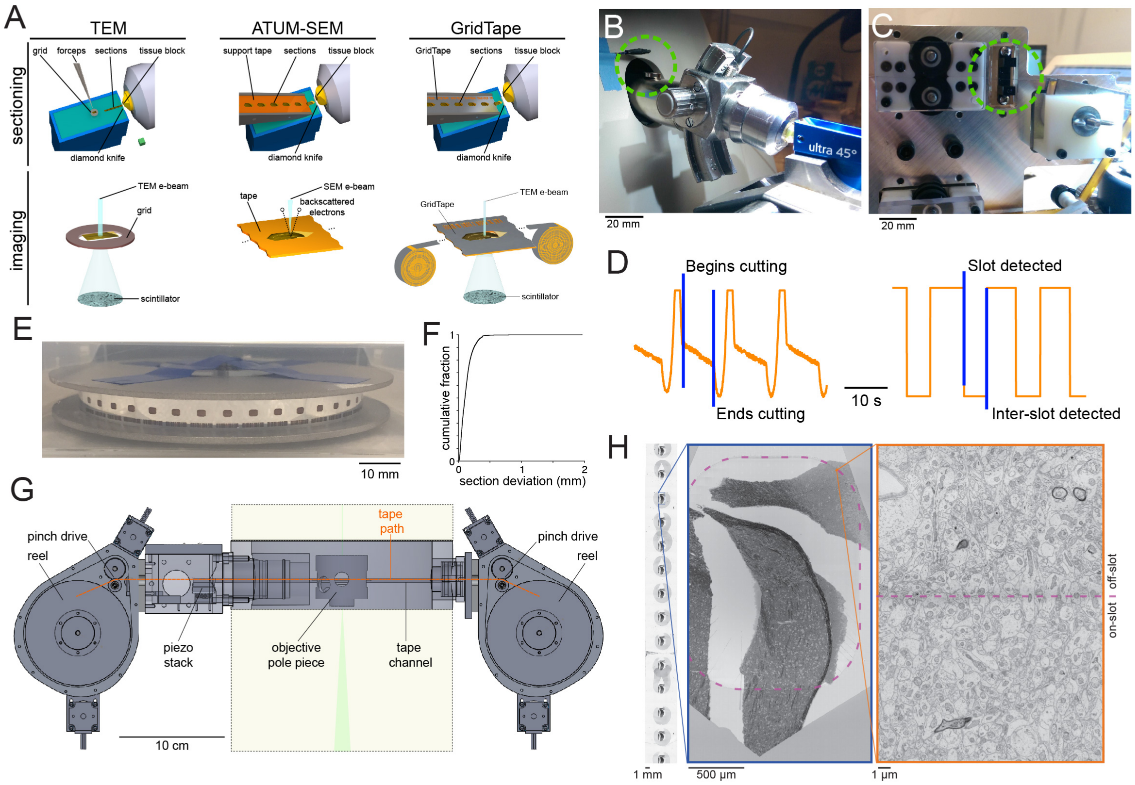

**Figure S1 (Related to Figure 1). The GridTape imaging platform.**

(A) Schematics of (left) manual serial-section collection and transmission electron microscopy (TEM) imaging; (middle) automated tape-collecting ultramicrotome (ATUM) collection and scanning electron microscopy (SEM) imaging, and (right) GridTape sectioning and TEM imaging. Ellipses denote continuation of tape. Bottom schematics not to scale.

(B) Photograph of the magnet and hall effect sensor (dashed circle) attached to the microtome cutting arm to measure cutting frequency.

(C) Photograph of the digital opto-interrupter (dashed circle) used to detect the slot frequency in the tape.

(D) The (left) analog signal from the hall effect sensor and (right) digital signal from the opto-interrupter are used to perform closed-loop, phase-locked collection of sections onto GridTape slots.

(E) Photograph of a reel of GridTape containing 4355 serial sections of an adult female *Drosophila* VNC.

(F) Cumulative distribution of section placement deviation (Euclidean distance from the average section position) across the VNC dataset.

(G) Schematic of the GridTape reel-to-reel stage at the level of the TEM objective pole-piece. Portions of the TEM column above and below in beige with dashed outlines. Electron beam in green (not to scale).

(H) Regions of sections not collected over slots can be imaged with SEM as in the ATUM-SEM approach. SEM images of (left) a stretch of GridTape carrying mouse thalamus sections, (middle) a single section over a slot, and (right) tissue collected over the edge of the slot. This approach was not required for the VNC dataset due to the low number of off-slot sections.

Scale box, 1 $mm^3$ (A, top), Scale bars, 20 mm (B-C), 10 mm (E), 1 mm (H, left), 500 µm (H, middle), 1 µm (H, right).

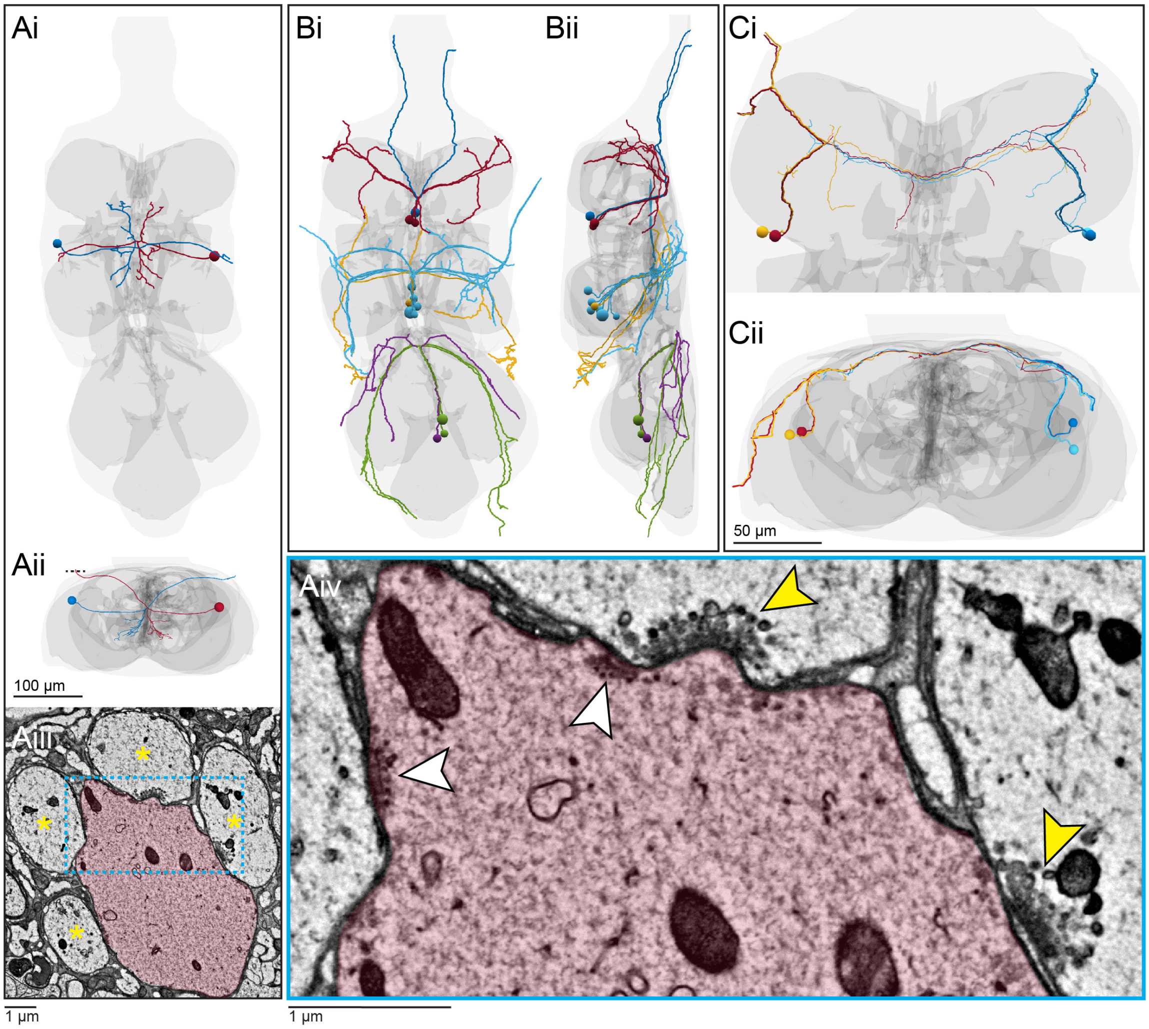

**Figure S2 (Related to Figure 2). Efferent neurons with unusual features.**
(A) The peripherally-synapsing interneuron (PSI). (i) Dorsal view. (ii) Anterior view. (iii) Cross-section through the posterior dorsal mesothoracic nerve at the location indicated by the dashed line in (ii), showing the PSI (red) fasciculated with four motor neurons (asterisks). (iv) Higher resolution view of the region indicated in (iii). The PSI makes synaptic outputs onto neighboring motor neurons (white arrowheads) as described previously (King and Wyman, 1980). Unexpectedly, some of the motor neurons appear to make synapses back onto the PSI (yellow arrowheads). These putative synapses are not observed in motor neurons in any other nerves.
(B) Dorsal unpaired median (DUM) neurons (Duch et al., 1999) with cell bodies organized into 3 clusters. (i) Dorsal view. (ii) Lateral view. Colors indicate target organ (see color key in Fig. 2A).
(C) A novel cell type, the "multinerve" neuron. (i) Dorsal view. (ii) Anterior view. These neurons have cell bodies lateral and posterior to each T1 neuromere, and they send projections out both the ipsilateral dorsal prothoracic nerve and ipsilateral prothoracic accessory nerve. Their branches are positioned on the dorsal-most surface of the VNC, including a contralateral projection. Multinerve neurons are unique to T1, as similar neurons were not found in T2 or T3.
Scale bars, 100 µm (Ai-ii, B), 1 µm (Aiii-iv), 50 µm (C).

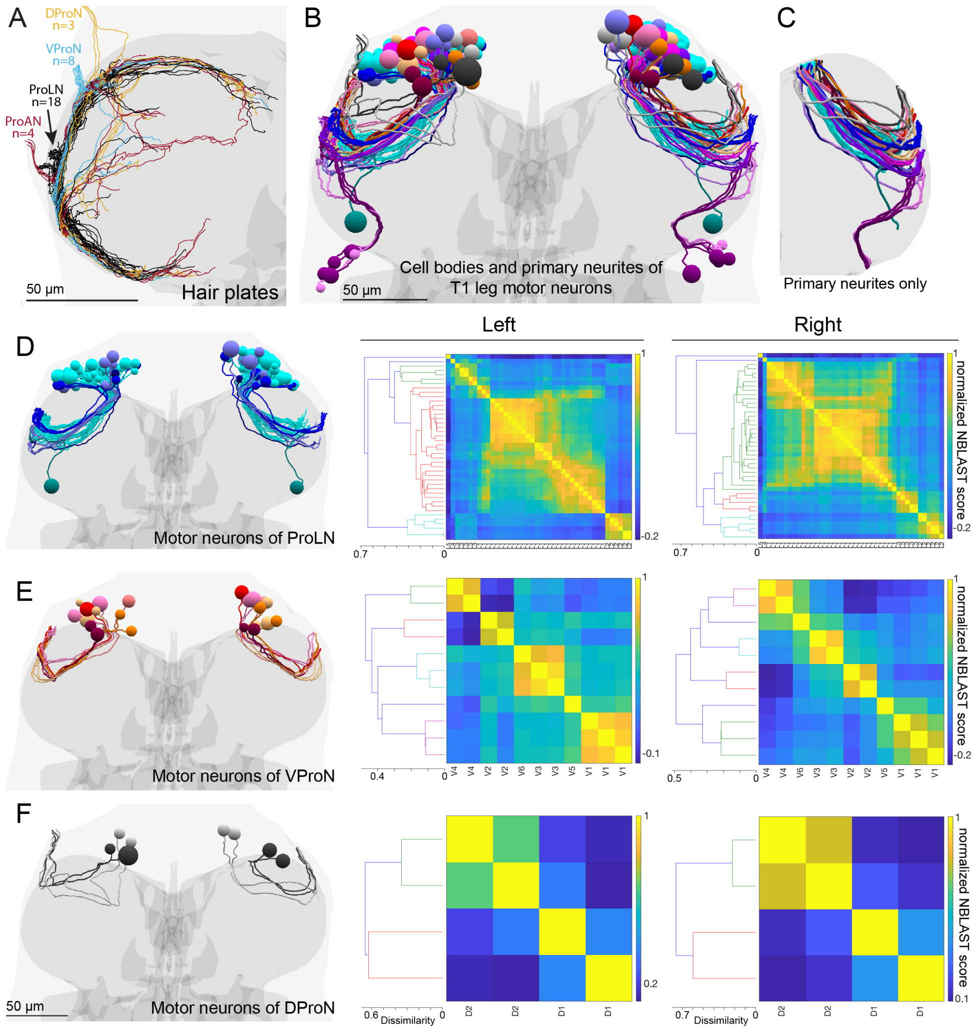

**Figure S3 (Related to Figures 3-4). Hair plate neurons and motor neuron bundles.**

(A) Organization of hair plate neuron projections in the VNC. Hair plate axons enter the T1 neuromere through four different nerves and branch to encircle the neuromere. The number of hair plate axons entering each nerve is indicated. Color coded by nerve.

(B) Reconstruction of the cell bodies and primary neurites of T1 leg motor neurons. Primary neurites travel through the neuromere in distinct bundles before leaving the VNC through one of four different peripheral nerves. The largest bundle (cyan) contains 29 primary neurites on the left side and 30 on the right side. The remaining 40 neurons per side had primary neurites organized into bundles containing between one and six members, with the left and right side bundles always containing the same numbers of members.

(C) Motor neurons were transformed into the VNC atlas coordinate system and pruned to exclude regions outside of the neuropil. The remaining portions included only the primary neurite and were used for calculation of all similarity scores in Fig. 4B and in (D-F) below.

(D-F) Rendering of primary neurites and cell bodies color coded by bundle (left) and hierarchical clustering dendrograms and matrices of NBLAST similarity scores (right), as in Fig. 4A-B but for motor neurons of different nerves.

(D) Prothoracic leg nerve (ProLN) motor neurons.

(E) Ventral prothoracic nerve (VProN) motor neurons.

(F) Dorsal prothoracic nerve (DProN) motor neurons.

Scale bars, 50 µm (A-E).

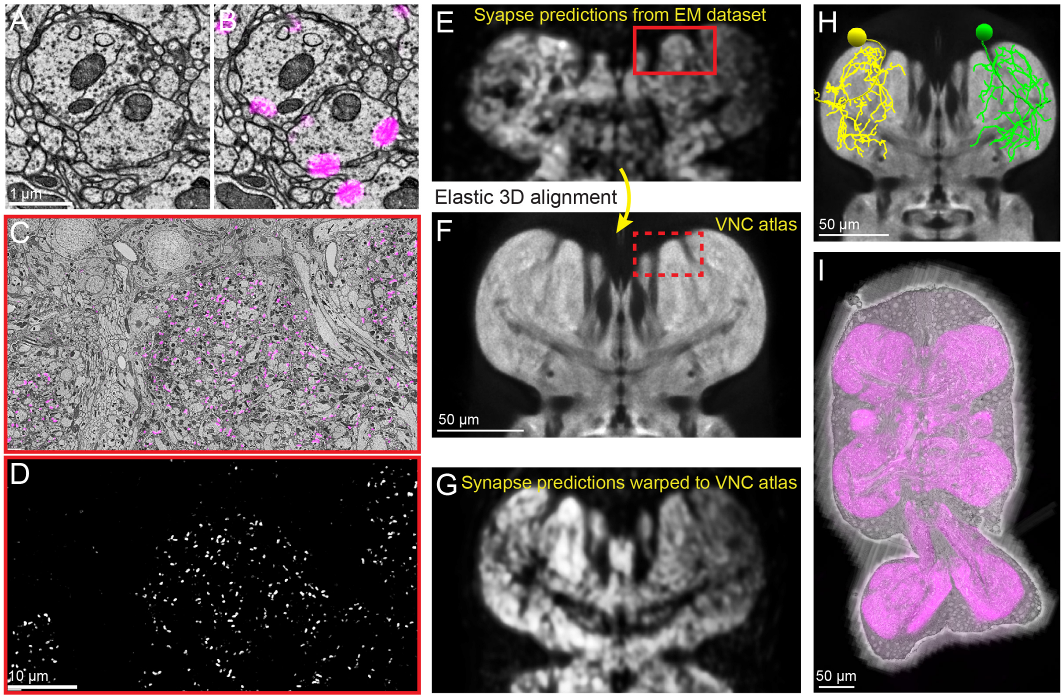

**Figure S4 (Related to Figures 4, 6). Automated synapse prediction and registration to a standard VNC atlas.**

(A-D) A convolutional neural network was used to predict presynaptic locations across the entire EM dataset.

(A) Raw EM data. Same data as Fig 1K.

(B) Raw EM data from (A) overlaid by predicted presynaptic locations (purple densities).

(C) A wider field of view showing raw EM data (greyscale) overlaid by predicted presynaptic locations (purple densities). Predicted synapses fall within the neuropil but are not present in the regions containing cell bodies or in the bundles of axons entering the neuropil.

(D) Predicted presynaptic locations from (C) but without EM data.

(E) The predicted synapse locations were 3D Gaussian blurred ($\sigma = 2$ µm) and downsampled to produce a synapse density map across the entire VNC EM dataset. Shown here is a slice through the predicted density map for the T1 neuromeres. Red box corresponds to the field of view from (C) and (D). The synapse-free locations are clearly distinguishable from synapse-containing regions.

(F) A slice through the female adult Drosophila VNC atlas, a synapse density map that serves as a reverence coordinate system. The location in the atlas corresponding to the boxed region in (E) is indicated (dashed red box).

(G) Synapse densities from the EM dataset underwent 3D elastic alignment to the VNC atlas. This enables data to be transformed between the EM dataset's coordinate system and atlas coordinates. Shown here is a slice through the synapses from the EM dataset transformed into the atlas coordinates. Notice the resemblance to (F). See also Video S4.

(H) Using the registration to the atlas, EM-reconstructed neurons can be transformed into the coordinate system of the atlas (yellow). This enables structural comparisons between EM-reconstructed neurons (see Fig. 4) and neurons imaged with light microscopy that have also been registered to the atlas (green, see Fig. 6).

(I) Using the registration, the VNC atlas was transformed into the EM dataset's coordinate system. Shown here is a slice of the transformed atlas (purple) overlaid on a slice of the EM dataset.

Scale bars, 1 µm (A-B), 10 µm (C-D), 50 µm, (E-I).

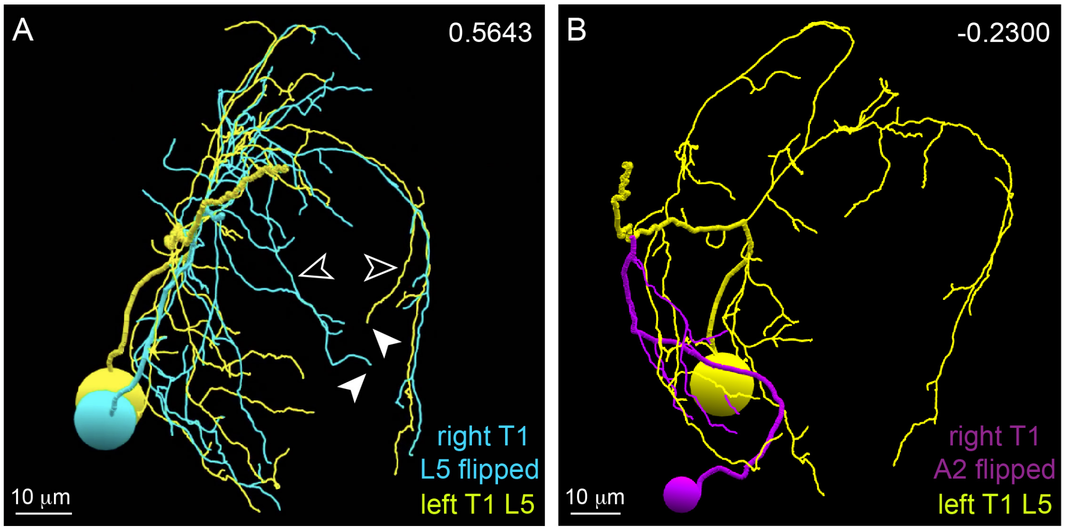

**Figure S5 (Related to Figures 4 and 6) Motor neuron similarity and variance.**
(A) Rendering of the left T1 L5 motor neuron and right T1 L5 motor neuron reflected across the midline. These motor neurons had cell bodies in a unique location (see Fig 3B) that allowed these two neurons to be unambiguously identified as a left-right homologous pair. NBLAST similarity score in the upper right corner. Scores between predicted homologous pairs typically ranged between 0.5 and 0.7, indicating that factors such as developmental variability between left and right copies of an identified neuron often results in NBLAST similarity scores in the 0.5 to 0.7 range. Note that while these neurons project to similar locations within the neuromere, their branches sometimes have different trajectories. Filled arrowheads point to example branch terminations with similar positions, with empty arrowheads pointing to their different paths. Differences like this may underlie why left-right homologous neuron pairs do not have a similarity score closer to 1.
(B) Rendering of the left T1 L5 motor neuron (perspective rotated relative to (A)) and the T1 leg motor neuron with the lowest NBLAST score.
Scale bars, 10 μm (A-B).

#### SUPPLEMENTAL TABLES

| | |
|---|---|
| JEOL 1200EX TEM | $ 150,000 |
| GridTape stage | $ 50,000 |
| Cameras (4) | $ 65,000 |
| Camera lenses (4) | $ 4,800 |
| Computers (5) | $ 12,500 |
| Vacuum parts | $ 5,000 |
| Leaded glass | $ 5,000 |
| Scaffolding | $ 5,000 |
| Total | $ 297,300 |

**Table S1. Related to Table 1. TEMCA-GT cost in current US$.**

#### SUPPLEMENTAL VIDEOS

**Video S1. Fly-through of the aligned VNC EM dataset.**
Zoomed-out (left) and zoomed-in (right) views of aligned image data. Section number in the upper left corner.

**Video S2. Reconstructed motor and sensory neurons in the VNC.**
Skeleton reconstructions of motor neurons (left) and sensory neurons (right) from the VNC EM dataset (gray surface rendering). Spheres represent neuronal cell bodies. These neurons connect the VNC to the front (red), middle (yellow), and hind (green) legs as well as the neck (blue), wings (cyan), and halteres (purple). Anterior is up and posterior is down.

**Video S3. Reconstructed sensory and motor neuron subtypes.**
Skeleton reconstructions of sensory and motor neuron axons in the left T1 neuromere color coded by type as in Fig. 3Ai, B: chordotonal (orange), campaniform sensilla (light orange), hair plates (light blue), bristles (purple). Primary neurites and cell bodies of motor neurons of the prothoracic leg nerve from different bundles are different shades of blue. Anterior is up and posterior is down.

**Video S4. Alignment of the EM dataset to a reference VNC atlas**
A convolutional neural network was used to predict the location of synapses in the VNC EM dataset. Those predictions were blurred and downsampled to produce a map of synapse densities across the VNC dataset (left). This enabled the EM dataset to be aligned to a reference VNC atlas (middle and right).

**Video S5. Gallery of left-right T1 motor neuron pairs.**
3D renderings of the 36 left-right pairs of T1 motor neurons (Fig 4D) after transformation into the VNC atlas coordinate system. Neurons are color coded by the bundle (see Figs. 4A and S3B-F for color code). Outline of the VNC atlas in grey. Spheres represent motor neuron cell bodies.

**Video S6. Bilateral campaniform sensilla (bCS) directly synapse onto a fast flexor motor neuron.**
bCS axons (red and blue traces) make output synapses (small red balls) directly onto specific motor neurons. bCS neurons synaptically target the highest NBLAST scoring match to the fast tibia flexor motor neuron (green, 81A07-Gal4), but not the best matching slow motor neuron (magenta, 35C09-Gal4).
